# Relabeling metabolic pathway data with groups to improve prediction outcomes

**DOI:** 10.1101/2020.08.21.260109

**Authors:** Abdur Rahman M. A. Basher, Steven J. Hallam

## Abstract

Metabolic pathway inference from genomic sequence information is an integral scientific problem with wide ranging applications in the life sciences. As sequencing throughput increases, scalable and performative methods for pathway prediction at different levels of genome complexity and completion become compulsory. In this paper, we present reMap (<u>re</u>labeling <u>m</u>etabolic pathway d<u>a</u>ta with grou<u>p</u>s) a simple, and yet, generic framework, that performs relabeling examples to a different set of labels, characterized as groups. A pathway group is comprised of a subset of statistically correlated pathways that can be further distributed between multiple pathway groups. This has important implications for pathway prediction, where a learning algorithm can revisit a pathway multiple times across groups to improve sensitivity. The relabeling process in reMap is achieved through an alternating feedback process. In the first feed-forward phase, a minimal subset of pathway groups is picked to label each example. In the second feed-backward phase, reMap’s internal parameters are updated to increase the accuracy of mapping examples to pathway groups. The resulting pathway group dataset is then be used to train a multi-label learning algorithm. reMap’s effectiveness was evaluated on metabolic pathway prediction where resulting performance metrics equaled or exceeded other prediction methods on organismal genomes with improved predictive performance.

## 1 Introduction

Biological systems operate on the basis of information flow between genomic DNA, RNA and proteins. Proteins catalyze most reactions producing metabolites. Reaction sequences are called pathways when they contribute to a coherent set of interactions driving metabolic flux within or between cells. Inferring metabolic pathways from genomic sequence information is a fundamental problem in studying biological systems with far-reaching implications for our capacity to perceive, evaluate and engineer cells at the individual, population, and community levels of biological organization [4, 8]. Over the past decade, the rise of next generation sequencing platforms has created a veritable tidal wave of organismal and multi-organismal genomes that must be assembled and annotated at scale without intensive manual curation. In response to this need, gene-centric and pathway-centric methods have been developed to reconstruct metabolic pathways from genomic sequence information at different levels of complexity and completion. The most common methods are gene-centric and involve mapping predicted protein coding sequences onto known pathways using a reference database (e.g. the Kyoto Encyclopedia of Genes and Genomes (KEGG) [6]. Alternative pathway-centric methods including PathoLogic [7] and MinPath [20] predict the presence of a given metabolic pathway based on heuristic or rule-based algorithms. While gene-centric methods are effective at producing parts list, they are unable to infer pathway presence or absence given a set of predicted protein coding sequences. Conversely, while pathway-centric methods infer pathway presence or absence given a set of predicted protein coding sequences, the development of reliable and flexible rule sets is both difficult and time consuming [19].

Machine learning methods aim to improve on heuristic or rule-based pathway inference through features engineering and algorithmic solutions to overcome noise and class imbalance. Basher and colleagues developed mlLGPR [11], a multi-label classification method that uses logistic regression and feature vectors inspired by the work of Dale and colleagues [3] to predict metabolic pathways from genomic sequence information at different levels of complexity and completion [11]. Recently, triUMPF ([12, 13]) was proposed to reconstruct metabolic pathways from organismal and mutli-organismal genomes. This method uses meta-level interactions among pathways and enzymes within a network to improve the accuracy of pathway predictions in terms of communities represented by a cluster of nodes (pathways and enzymes). Despite triUMPF’s predictive gains, its sensitivity scores on pathway datasets left extensive room for improvement. Here, we present reMap that relabels each example with a new label set called “pathway group” or “group” forming a pathway group dataset which then can be employed by a suitable pathway prediction algorithm (e.g. leADS [14]) to improve prediction results.

A subset of pathways in multiple organisms may be statistically correlated and this subset constitutes a group. Thus, the presence of a pathway entails the presence of a set of other correlated pathways. reMap performs an iterative procedure to group statistically related pathways into a set of “pathway groups” using a correlation model (CTM, SOAP, and SPREAT see Appx. Section B). reMap then annotates organismal genomes with relevant groups. Pathways in these groups are correlated and allowed to be inter-mixed across groups with different proportions, resulting in an overlapping subset of groups over a subset of pathways (i.e., non-disjoint). This has important implications for pathway prediction, where a learning algorithm can revisit a pathway multiple times across groups to improve sensitivity. Unlike mlLGPR [11] and triUMPF (Fig. 1a), group based pathway prediction requires two consecutive parts. First, a set of pathway groups are inferred. In the second, pathways in these groups are predicted (Fig. 1b).

**Fig. 1:**
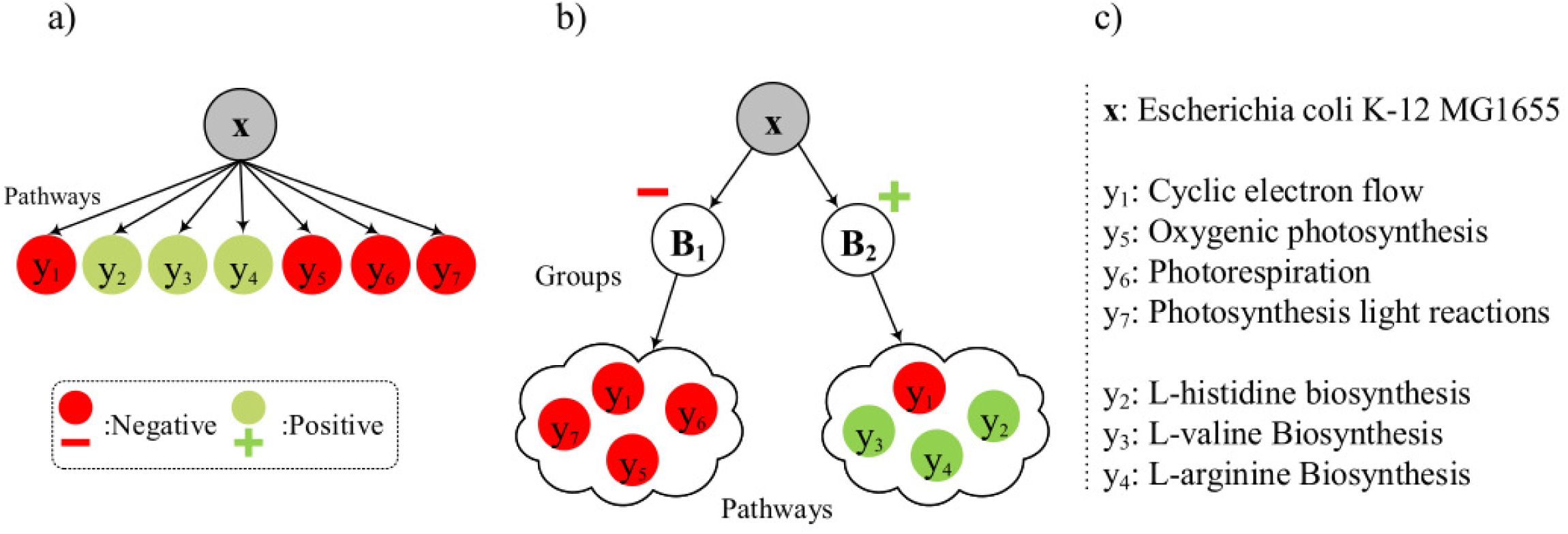
Traditional vs proposed group-based pathway prediction methods. In the traditional method (a) pathways (i.e., *y*^1−7^) are predicted for Escherichia coli K-12 MG1655, denoted by **x**, without considering any grouping of pathways. In contrast, the group-based pathway prediction method (b) uses a two step process. First, it predicts a set of positive groups (i.e., **B**_2_), then the pathways within these groups are predicted (depicted as a cloud glyph and true pathways are green colored). The description of symbols is provided in subfigure (c).

reMap’s pathway grouping performance was compared with other methods including MinPath, PathoLogic, and mlLGPR on a set of Tier 1 (T1) pathway genome databases (PGDBs), low complexity microbial communities including symbiont genomes encoding distributed metabolic pathways for amino acid biosynthesis [15], genomes used in the Critical Assessment of Metagenome Interpretation (CAMI) initiative [16], and whole-genome shotgun sequences from the Hawaii Ocean Time Series (HOTS) [17] following the genomic information hierarchy benchmarks initially developed for mlLGPR enabling more robust comparison between pathway prediction methods [11].

## 2 Method

In this section, we provide a general description of the reMap method, presented in Fig. 2. reMap is trained in two phases using an alternating feedback process: i)- feed-forward in Figs 2(b-d), consisting of three components: 1)- constructing pathway group, 2)- building group centroid, 3)- mapping examples to groups; and ii)- feed-backward to update reMap’s parameters in Fig. 2(f). After training is accomplished, a pathway group dataset is produced that can be used to predict metabolic pathways from a newly sequenced genome in Figs 2(g-h). Below, we discuss these two phases while the analytical expressions of reMap are explained in Appx. Section A.

**Fig. 2:**
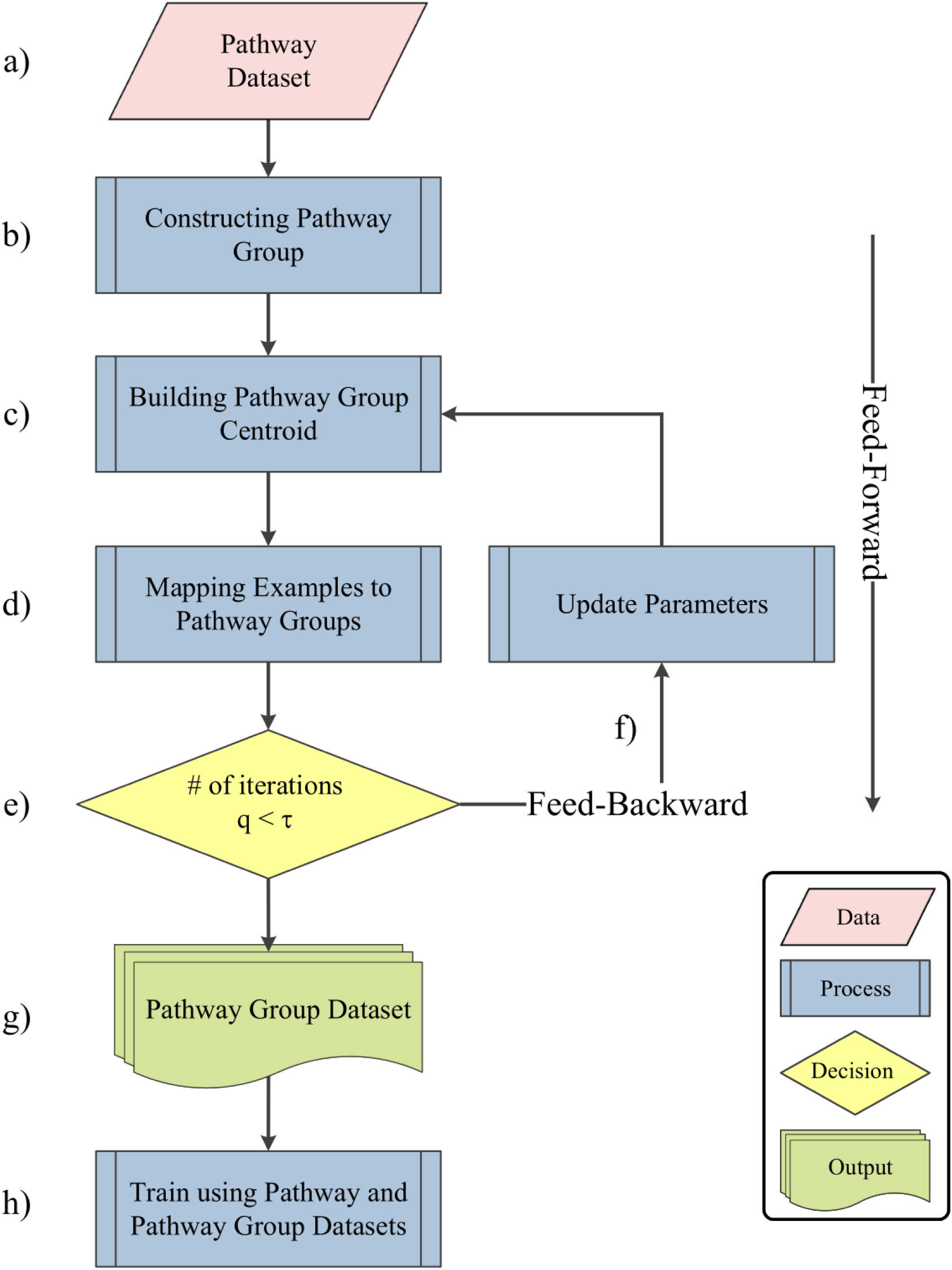
A workflow diagram for reMAP. The relabeling process in reMap is achieved through an alternating feedback process. The feed-forward phase is composed of three components: (b) pathway group construction to build correlated pathway groups from pathway data (a), (c) building group centroid to estimate centroids of groups, and (d) mapping examples to groups. The feed-backward phase (f) optimizes reMap’s parameters to increase accuracy of mapping examples to groups. The process is repeated *τ* (∈ 𝕫_*>*1_) times. If the current iteration *q* (∈ 𝕫_*>*1_) reaches the desired number of rounds *τ*, the training is terminated (e) and the pathway group dataset is produced (g) which can be used as inputs to a pathway inference algorithm (e.g. leADS [14]) to predict a set of pathways from a newly sequenced genome (h).

### 2.1 Feed-Forward Phase

During this stage, each example in a given pathway data (Appx. Def. 1) is annotated with a subset of pathway groups in three consecutive steps:

#### Constructing Pathway Group

In this step, pathways are partitioned into non-disjoint *b* (∈ 𝕫_≥1_) groups using any correlation models defined in Appx. Section B. These models are equipped to provide us with a group correlation matrix and a pathway distribution over groups, denoted by *Φ* ∈ ℝ^*b×t*^, where *t* corresponds to the total number of distinct pathways. Each entry *Φ*_*i,j*_ corresponds to the probability of assigning a pathway *j* to the group *i*. For each group in *Φ*, we retain the top *k* (∈ 𝕫_≥1_) pathways based on the probability scores. The trimmed *Φ* serves as an input to constructing centroids in the next step.

Modeling pathway distribution and group correlation in this way are motivated by two key intuitions. First, organisms encoding similar pathways may share similar groups resulting in shared statistical properties for those organisms. Second, frequently occurring pathways in multiple organisms imply a similar relative contribution to a group.

#### Building Group Centroid

Having obtained a set of groups, reMap determined the relative contribution of each pathway to its associated group’s centroid in the Euclidean space. Estimating centroids requires representing pathways and groups as vectors of real numbers. For this, we apply pathway2vec [10] to obtain pathway features. Then, the centroid of a group, say *s*, is computed as:

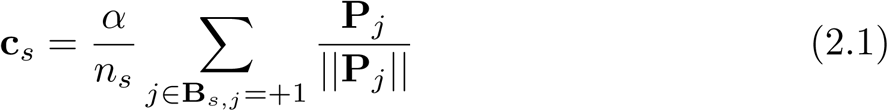

where **B**_*s*_ ∈ {−1, +1}^*t*^is the group *s* obtained from the trimmed *Φ*_*s*_ after transforming it to {−1, +1}^*t*^. **c**_*s*_ corresponds the centroid of the group *s*, **P** ∈ ℝ^*t×m*^ is a pathway representation matrix obtained from pathway2vec, *n*_*s*_ is the number of pathways (|{**B**_*s,j*_ = +1, ∀*j* ∈ *t*}|) in group *s*, || · || is the length of a feature vector, and *α* (∈ ℝ_*>*0_) is a hyper-parameter determined by empirical analysis (16 in this work). The proposed Eq. 2.1 is based on the intuition that pathways associated with a group are semantically “close enough” to the center of the corresponding group, and the overlapping pathways among groups exhibit similar semantics with their associated groups. In addition to determining centroids, reMap also estimates a maximum number of expected groups to be annotated for a given example, indexed by *i*, using the cosine similarity metric [9]:

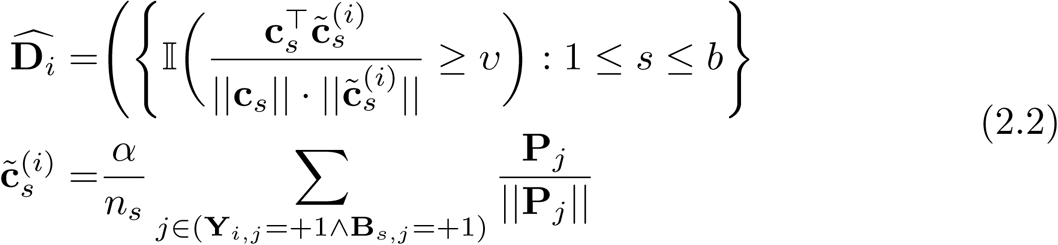

where 𝕀(.) is an indicator function that results in either +1 or 1 depending on a user-defined threshold *υ* (∈ ℝ_*>*0_). **Y**_*i*_ ∈ {1, +1}^*t*^ corresponds to pathways either present or absent for the *i*th example, indicated by +1 and 1, respectively. 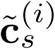 represents the centroid of the group *s* calculated based on pathways that are associated with the group *s* and are present in *i*th example. ñ_*s*_ is the number of pathways (|{**Y**_*i,j*_ = +1 ∧ **B**_*s,j*_ = +1, ∀*j* ∈ *t*}|) in group *s*. 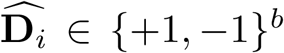 is a pre-optimized set of groups labelled for the *i*th example that will be used in the mapping step.

#### Mapping Pathways to Pathway Groups

This step maps an example to pathway groups, resulting in an optimized pathway group dataset 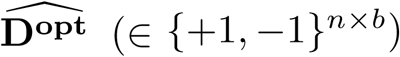. Formally, let us denote a set of groups that are picked to label an example by 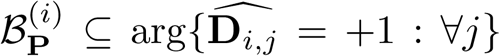 while the remaining unpicked groups is denoted by 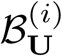, where 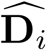 is obtained using Eq. 2.2. Both sets of groups are stored in 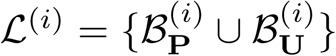. Then, reMap performs mapping in an iterative way, mirroring sequential learning and prediction strategy [18], where for each *i*th example, a group **B**_*j*_ at round *q* is either: i)-added to *ℒ*^(*i*)^, indicated by 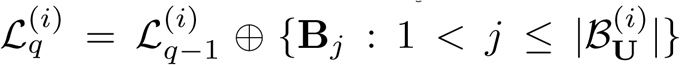;or ii)- removed from the set of selected groups, represented by 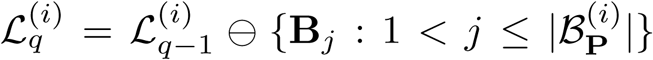. More specifically, at each iteration *q*, reMap estimates the probability of an example, given the selected groups that are obtained from the previous round *q* − 1, using the threshold closeness (TC) metric [2] as:

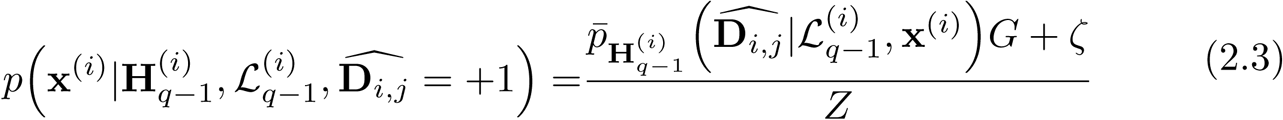

where **x**^(*i*)^ ∈ ℝ^*r*^ and *r* is the total number of enzymes, 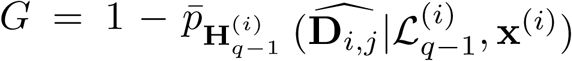 and 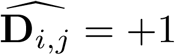 if the group **B**_*j*_ is tagged with the *i*th example.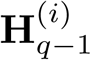 represents the history of prediction probability storing all 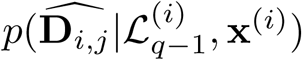 before the current iteration *q* while 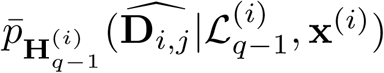 is the average probability of classifying **x**^(*i*)^ to the group **B**_*j*_ over values in 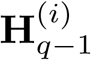.The term *ζ*(∈ ℝ_*>*0_) is a smoothness constant and *Z* is a normalization constant. Note that TC is a class conditional probability density function that encourages correct class probability to be close to the true unknown decision boundary. Hence, this step will ensure the correct latent group to be assigned to the *i*th example. The parameter 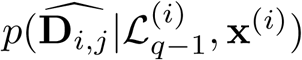 can be estimated using Appx. Eq. A.4. Afterwards, ℒ^*(i)*^ will be updated either by adding or removing groups from a previous iteration. More details about this step is provided in Appx. Section A.1.

### 2.2 Feed-Backward Phase

During this phase, reMap updates its internal parameters by enforcing four constraints: i)- similarity between groups and associated pathways; ii)- weights of pathways, in a group, should be close to each other; iii)- examples sharing similar pathways should share similar representations; and iv)- all reMap’s parameters should not be too large or too small. These four constraints are important to allow smooth updates and mapping operations. More details are provided in Appx. Section A.2.

### 2.3 Closing the loop

The two phases are repeated for all examples in a given pathway data, until a predefined number of rounds *τ* (∈ 𝕫_*>*1_) is reached. At the end, a pathway group dataset is produced which consists of *n* examples with the assigned groups, i.e.,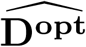. After training is accomplished, a pathway group dataset is produced that can be used to predict metabolic pathways from a newly sequenced genome using an ML prediction method such as leADS [14].

## 3 Experiments

We evaluated reMap’s performance on diverse pathway datasets traversing the genomic information hierarchy [11]: i)- T1 golden consisting of EcoCyc, HumanCyc, AraCyc, YeastCyc, LeishCyc, and TrypanoCyc; ii)- BioCyc (v20.5 T2 & 3) [1]; iii)- *Symbionts* genomes of *Moranella* (GenBank NC-015735) and *Tremblaya* (GenBank NC-015736) encoding distributed metabolic pathways for 9 amino acid biosynthesis [15]; iv)- Critical Assessment of Metagenome Interpretation (CAMI) dataset composed of 40 genomes [16]; and v)- whole genome shotgun sequences from the Hawaii Ocean Time Series (HOTS) at 25m, 75m, 110m (sunlit) and 500m (dark) ocean depth intervals [17]. Information about these datasets is presented in Appx. Section C.1.

Two experiments were conducted: i)- assessing the history probability and ii)- metabolic pathway prediction. The goal of the former test is to analyze the accumulated probability stored in **H** during the mapping process in the feed-forward phase for golden T1 datasets. We expect that few groups containing statistically related pathways will be annotated for T1 golden data. The metabolic pathway prediction test is followed to verify the quality of pathway groups for T1 golden, symbionts, CAMI, and HOTS data. For comparative analysis, reMap’s performance on T1 golden datasets was compared to four pathway prediction methods: i)- MinPath version 1.2 [20], an integer programming based algorithm; ii)- PathoLogic version 21 [7], a symbolic approach that uses a set of manually curated rules to predict pathways; iii)- mlLGPR [11], a supervised multi-label classification and rich feature information algorithm, and iv)- triUMPF [12, 13], a non-negative matrix factorization and community detection based algorithm. Four metrics were used to report the performance of all pathway predictors for golden T1 and CAMI data: *average precision, average recall, average F1 score (F1)*, and *Hamming loss* as described in [11]. In addition, reMap’s performance was compared to PathoLogic, mlLGPR, and triUMPF on mealybug symbionts and HOTS multi-organismal datasets. To construct pathway groups, we employed the correlated model SOAP using *b* = 200 groups.

reMap was written in Python v3 and is available under the GNU license at https://github.com/hallamlab/reMap. Unless otherwise specified all tests were conducted on a Linux server using 10 cores of Intel Xeon CPU E5-2650. For full experimental settings and additional tests, see Appx. Sections C and D.

### 3.1 Accumulated History Probability Analysis

Fig. 3a shows **H** during the annotation process for the T1 golden data over 10 iterations. In the beginning, reMap attempts to select the maximum number of groups that may exist for each example. However, with progressive updates and calibration of parameters, reMap rectifies groups assignments where it picks fewer relevant groups for each example. As an example, after the 10th round, Escherichia coli K-12 MG1655 was tagged with only 33 groups (Fig. 3b) and 18 of these groups contain amino acid biosynthesis pathways. Fig. 3c shows 7 of these 18 pathway groups (Appx. Table 5). Pathways in these 7 groups are statistically related (Appx. Table 6), and are observed to be distributed across groups reflected by the thickness of edges in Fig. 3c. For example, *L-alanine biosynthesis II* pathway is present in groups indexed by 16 and 152. Similarly, for the pathway *L-glutamate biosynthesis III* which is represented in groups indexed by 13 and 140. This mixture of pathway representation over groups increases the chance of a pathway inference algorithm (e.g. leADS [14]) to revisit a true positive pathway multiple times across groups which may result in improved predictions as reported in the next section. This experiment shows that reMap is able to capture statistically relevant pathways and map related groups to each example with a high degree of correlation.

**Fig. 3:**
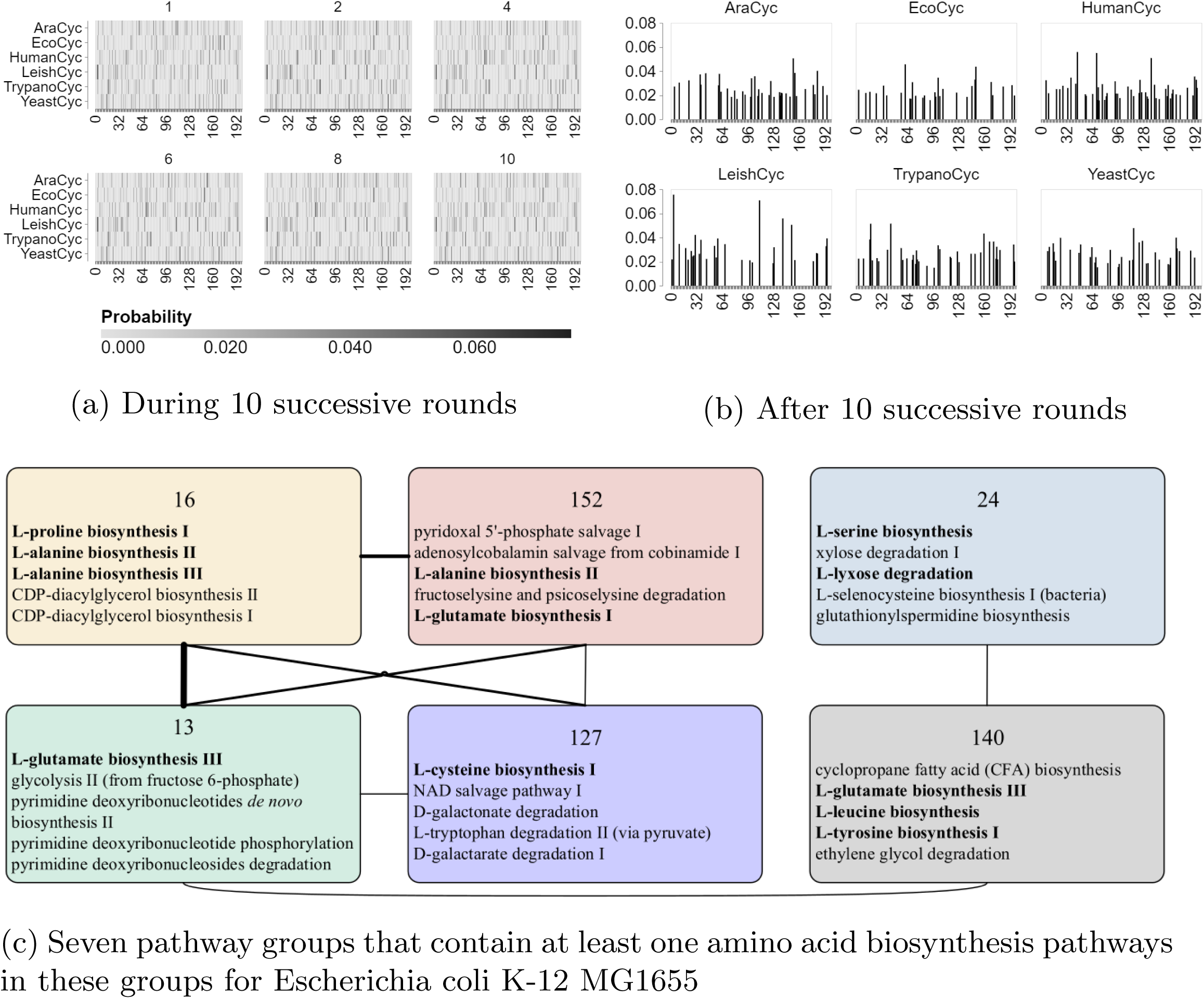
Fig. 3a illustrates the history probability **H** during annotation of T1 golden data over 10 successive rounds while Fig. 3b shows the results after 10th round. Darker colors indicate higher probabilities of assigning groups to the corresponding data. Fig. 3c shows six pathway groups and their correlations for Escherichia coli K-12 MG1655. Numbers at top boxes correspond to group indices. Edge thickness reflects the degree of associations between groups. Boldface text represent amino acid biosynthesis pathways.

### 3.2 Metabolic Pathway Prediction

#### T1 Golden data

Table 1 shows that reMap+SOAP achieved competitive performance against the other methods in terms of average F1 score with optimal performance on EcoCyc (0.8336). However, it under-performed on AraCyc, YeastCyc, and LeishCyc, yielding average F1 scores of 0.4764, 0.4914, and 0.4144, respectively. Since reMap+SOAP was trained using BioCyc containing less than 1460 trainable pathways, pathways outside the training set will be neglected.

**Table 1:**
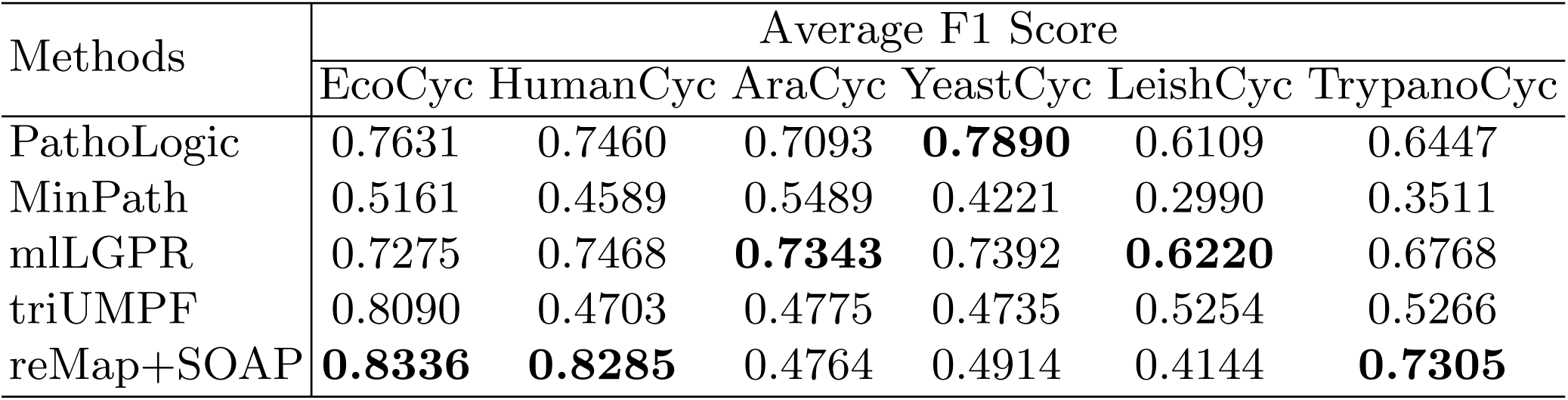
Average F1 score of each comparing algorithm on 6 golden T1 data. Bold text suggests the best performance in each column.

#### Symbionts data

The goal of this test is to evaluate reMap+SOAP performance on distributed metabolic pathways that emerge as a result of interactions between two or more organisms. We used the reduced genomes of *Moranella* and *Tremblaya* [15] as an established model for benchmarking. The two symbiont genomes in combination encode 9 intact amino acids biosynthesis pathways. All four pathway predictors were used to predict pathways on individual symbiont genomes and a composite genome consisting of both. While reMap+SOAP, triUMPF and PathoLogic predicted 6 of the expected amino acid biosynthesis pathways on the composite genome, mlLGPR was able to predict 8 pathways (Fig. 4). We excluded phenylalanine biosynthesis (*L-phenylalanine biosynthesis I*) pathway from analysis because the associated genes were reported to be missing after initial gene prediction. Four predictors identified false positives for individual symbiont genomes in *Moranella* and *Tremblaya* although the pathway coverage information for both genomes was reduced in relation to the composite genome (Appx. Fig. 9).

**Fig. 4:**
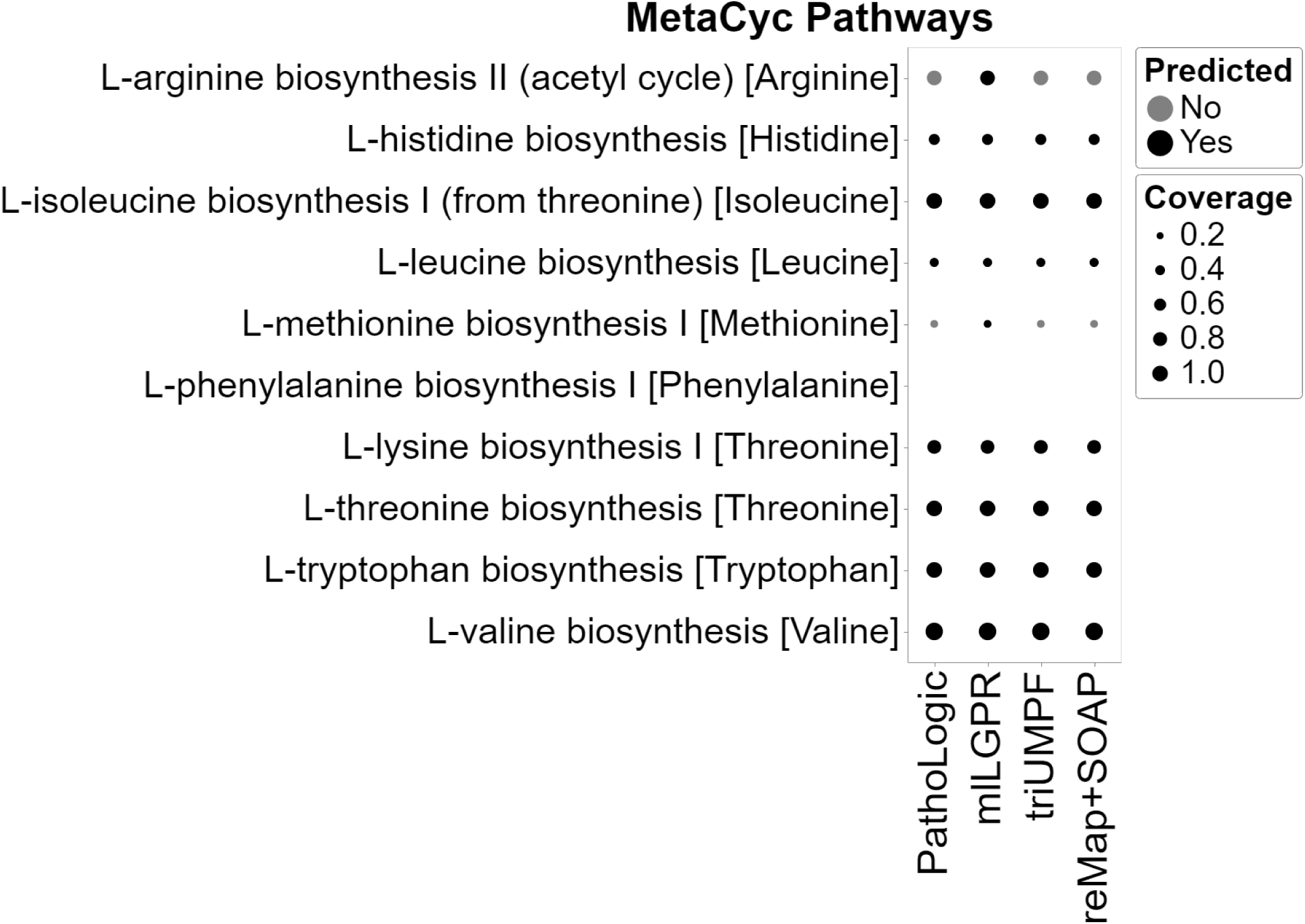
Comparative study of predicted pathways for the composite genome between PathoLogic, mlLGPR, triUMPF, and reMap+SOAP. Black circles indicate predicted pathways by the associated models while grey circles indicate pathways that were not recovered by models. The size of circles corresponds the pathway coverage information.

#### CAMI and HOTS data

For CAMI low complexity data [16], reMap+SOAP exceeded mlLGPR and triUMPF, achieving an average F1 score of 0.6125 in compare to 0.4866 for mlLGPR and 0.5864 for triUMPF (Table 2). For HOTS data [17], triUMPF, mlLGPR, and PathoLogic predicted a total of 58, 62, and 54 pathways, respectively, while reMap+SOAP inferred 67 pathways (see Appx. Section D.3) from a subset of 180 selected water column pathways [5]. None of the algorithms were able to predict pathways for *photosynthesis light reaction* and *pyruvate fermentation to (S)-acetoin* despite the abundance of these pathways in the water column. Absence of specific EC numbers associated with each pathway likely contributed to their absence using rule-based or ML prediction algorithms. Results from this experiment indicates that the proposed pathway group based approach, in particular reMap+SOAP increases pathway prediction performance relative to other methods used in isolation.

**Table 2:**
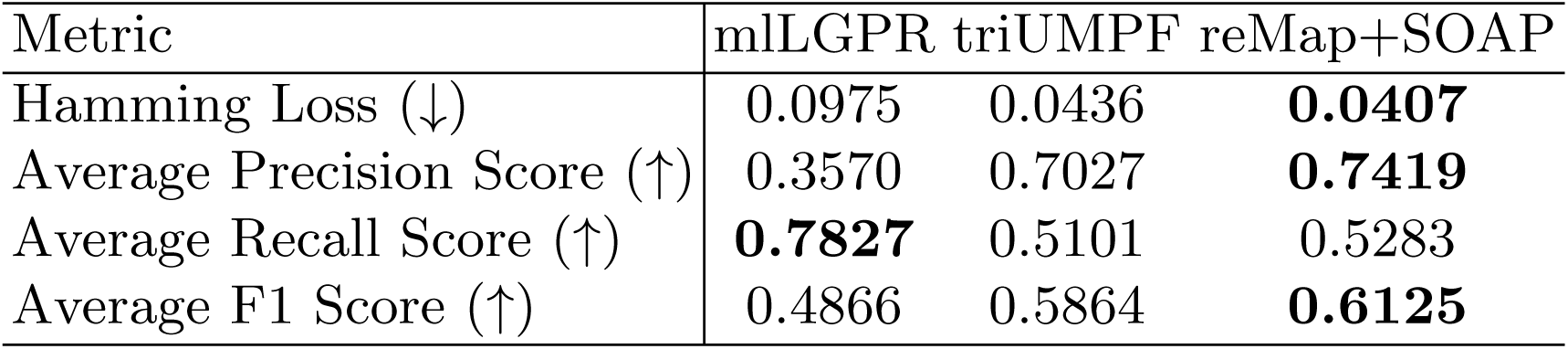
Predictive performance of mlLGPR, triUMPF, and reMap+SOAP on CAMI low complexity data. For each performance metric, ‘↓’ indicates the smaller score is better while ‘↑’ indicates the higher score is better.

## 4 Conclusion

In this paper, we demonstrated that iteratively mapping examples to groups e.g. relabeling, using reMap increased pathway prediction performance. The reMAP method is based on the intuition that organisms sharing a similar set of metabolic pathways may exhibit similar higher-level structures or groups. The relabeling process in reMap is achieved through an alternating feedback process. In the first feed-forward phase, a minimal subset of pathway groups is picked to label each example. In the second feed-backward phase, reMap’s internal parameters are updated to increase the accuracy of mapping examples to pathway groups. After training reMap, a pathway group dataset is produced that can be used to predict metabolic pathways for a newly sequenced genome.

We evaluated reMap’s performance for the pathway prediction task using a corpus of experimental datasets and compared results to other prediction methods including PathoLogic, MinPath, mlLGPR, and triUMPF. Overall, reMap showed promising results in boosting prediction performance over ML-based algorithms, such as mlLGPR and triUMPF. During benchmarking, we realized that reMap brings more frequent and sometimes irrelevant pathways, resulting in a significant performance loss on some T1 golden data, such as AraCyc. A possible treatment would be adding constraints in the form of associations among enzymes and pathways as applied in triUMPF. However, this may lead to sensitivity loss [14]. Another approach is to combine both graph-based and group-based strategies to predict pathways. Future development efforts will explore this dual approach to improve pathway prediction performance with emphasis on multi-organismal genomes encoding distributed metabolic processes.

## Appendix

The appendix is divided into four parts: i)- the reMap framework (Section A), ii)- descriptions about correlated models (Section B), iii)- experimental settings (Section C), and iv)- empirical analysis (parameter sensitivity, history probability analysis, and metabolic pathway prediction) (Section D).

### A The reMap Method

In this section, we provide important notations and definitions that will be used throughout the paper followed by a formal description of the research problem. All vectors are assumed to be column vectors and are represented by boldface lowercase letters (e.g., **x**) while matrices are encoded by boldface uppercase letters (e.g., **X**). The **X**_*i*_ matrix indicates the *i*-th row of **X** and **X**_*i,j*_ denotes the cell entry (*i, j*) of **X**. A subscript character to a vector, **x**_*i*_, denotes an *i*-th cell of **x**. Occasional superscript, **X**^(*i*)^, suggests an index to a example or current epoch during the learning period. The sets are characterized by calligraphic letters (e.g., *E*) while we use the notation |.| to denote the cardinality of a given set. With these notations, we introduce the problem examined in this paper, starting with a multi-label pathway dataset definition.

#### Definition 1. Multi-label Pathway Dataset [11]

*A general form of pathway dataset is characterized by 𝒮* = {(**x**^(*i*)^, **y**^(*i*)^) : 1 *< i* ⩽ *n*} *consisting of n examples, where* **x**^(*i*)^ *is a vector indicating the abundance information corresponding the enzymatic reactions. An enzymatic reaction, in turn, is denoted by e, which is an element of a set of enzymatic reactions E* ={*e*_1_, *e*_2_, …, *e*_*r*_}, *having r possible reactions, hence, the vector size* **x**^(*i*)^ *is r. The abundance of an enzymatic reaction for an example i, say* 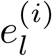, *is defined as* 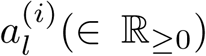. *The class labels* 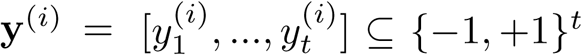 *is a pathway label vector of size t that represents the total number of pathways, which themselves are derived from a set of universal metabolic pathway 𝒴 The entry* +1 *(or* −1*) indicates presence (or absence) of a pathway corresponding the example i. The matrix form of* **x** *and* **y** *are symbolized as* **X** *and* **Y**, *respectively*.

Both *E* and *𝒴* are extracted from reliable knowledge-bases (e.g. KEGG [6] and MetaCyc [26]). In this paper, we adopt MetaCyc. Moving on, we define the term *pathway group set*.

#### Definition 2. Pathway Group Set

*Denote 𝔅* = {**B**_1_, **B**_2_, …, **B**_*b*_} *a set with b pathway groups, where each group* **B**_*c*_ ∈ {−1, +1}^*t*^*is presumed to contain a subset of correlated pathways, i*.*e*., *𝒴*_*c*_ ⊆ *𝒴, and t is the number of pathways in Def. 1*. *The presence or absence of a pathway in a group c is indicated by* +1 *or* −1, *respectively. The matrix representation of 𝔅 is* **B** ∈ {−1, +1}^*b×t*^.

Pathway groups are also assumed to be correlated, i.e, non-disjoint, and can be modeled by a Gaussian covariance matrix, denoted by *Σ* ∈ ℝ^*b×b*^. Each entry *s*_*i,j*_ in *Σ* characterizes the *i*-th group association with *j*-th group, where a larger score indicates both groups are highly correlated. As a result of correlation, we define the following two terminologies: *pathway group feature vector* and *pathway group’s neighbor*.

#### Definition 3. Group-Example Feature Vector

*The pathway group feature vector for the ithe example is indicated by* **d**^(*i*)^ ∈ {−1, +1}^*b*^, *where* 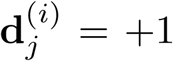 *iff the group j is observed for the example i and* **d**_*j*_ = −1 *otherwise. The matrix form in represented as* **D** ∈ {−1, +1}^*n×b*^.

An example of feature vectors for groups is illustrated in Fig. 5, where 2- dimensional feature vectors for groups encode presence or absence of two groups **B**_1_ and **B**_2_, given a set of 6 pathways and pathway-group association information, depicted as a cloud glyph.

**Fig. 5:**
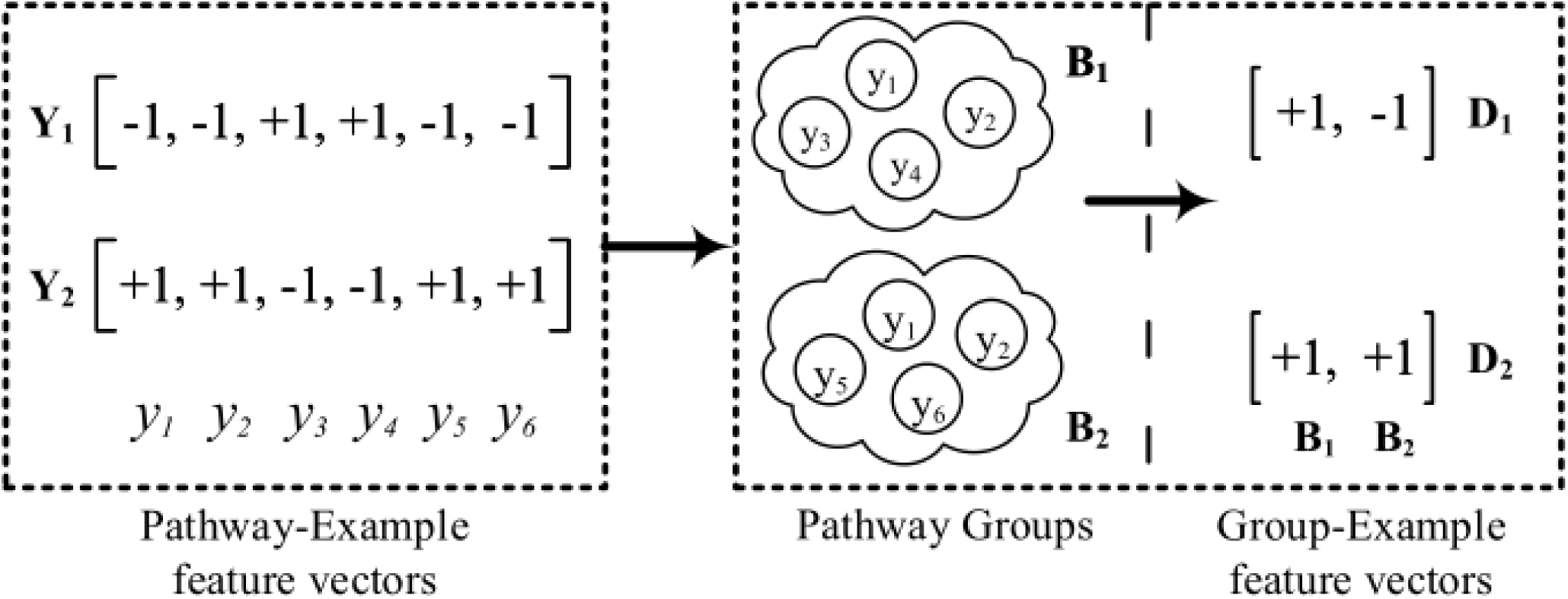
An example of feature vectors for groups. The subfigure in the left represents the feature vector for six pathways corresponding to two examples. The right subfigure indicates two groups, **B**_1_ and **B**_2_, and their features for the same two examples. The first example, **D**_1_, suggests that only **B**_1_ is present because the corresponding pathways *y*_3_ and *y*_4_ are present, while the pathway group feature vector for the second example, **D**_2_, suggests that both groups are present.

#### Definition 4. Pathway Groups Neighbors

*A group* **B**_*c*_ ∈ *𝔅 is said to be a neighbor to another pathway group* **B**_*j*_ ∈ *𝔅 s*.*t. c*≠ *j, if there exits an intersected pathway l in both groups, i*.*e*., **B**_*c,l*_ ∧ **B**_*j,l*_ = +1.

With the above definitions, we formulate the problem in this work.

*Problem 1*. Given a set of groups *𝔅* and a multi-label pathway dataset *𝒮*, the goal is to learn an optimum relabeling function *h*^**g**^ : → {+1, −1}^*b*^, such that leveraging groups to **X** incurs a high predictive score for the downstream pathway prediction task.

#### Algorithm 1: GroupCentroid(**P**, *𝔅, α*)

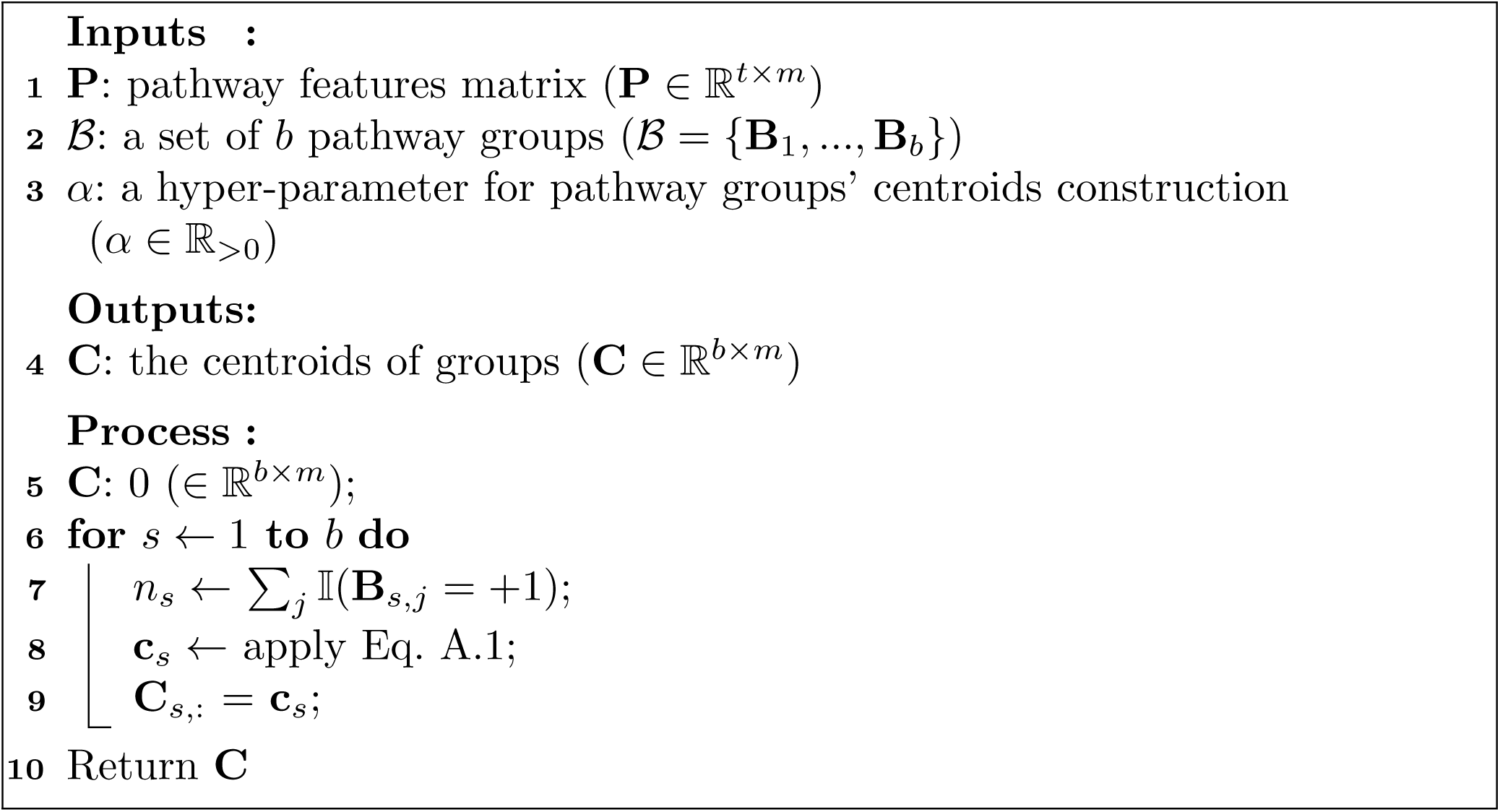

Fig. 1 illustrates the benefit of incorporating groups for multi-label pathway classification (right panel). Here, a dataset consists of two groups, each consists of a set of 4 correlated pathways. To determine positive pathways (*y*_2_, *y*_3_, and *y*_4_) given **X**_*i*_, we first predict the relevant group, indicated by +, then classify pathways within that pathway group. In contrast, the traditional multi-label classification approaches (left figure), mostly based on *binary relevance* technique, proceeds on predicting multiple pathway labels for **X**_*i*_. Hence, the proposed method will reduce computational complexity for pathway prediction.

Mapping a multi-label pathway dataset *𝒮* to groups will result in another dataset, i.e., *𝒮*_*group*_.

#### Definition 5. Multi-label Pathway Group Dataset

*A group dataset is represented by 𝒮*_*group*_ = {(**x**^(*i*)^, **d**^(*i*)^) : 1 *< i* ⩽ *n*} *consisting of n examples* 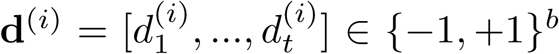 *is a pathway group label vector of size b. Each element of* **d**^(*i*)^ *indicates the presence/absence of the associated pathway group that is inherited from the set 𝔅 in Def. 2*.

Now, we outline the reMap method (depicted in Fig. 2), which alternates between the following two phases: i)- feed-forward in Figs 2(b-d), consisting of three components: 1)- constructing pathway group, 2)- building group centroid, 3)- mapping examples to groups; and ii)- feed-backward to update reMap’s parameters in Fig. 2(f). After training is accomplished, a pathway group dataset is produced that can used to predict metabolic pathways from a newly sequenced genome in Figs 2(g-h).

#### A.1 Feed-Forward Phase

During this phase, a minimal subset of groups is picked to annotate each example in a given pathway data (Def. 1) in three consecutive steps:

##### Algorithm 2: MaxGroups(*n*, **X, Y, P**,*𝔅*, **C**, *v*)

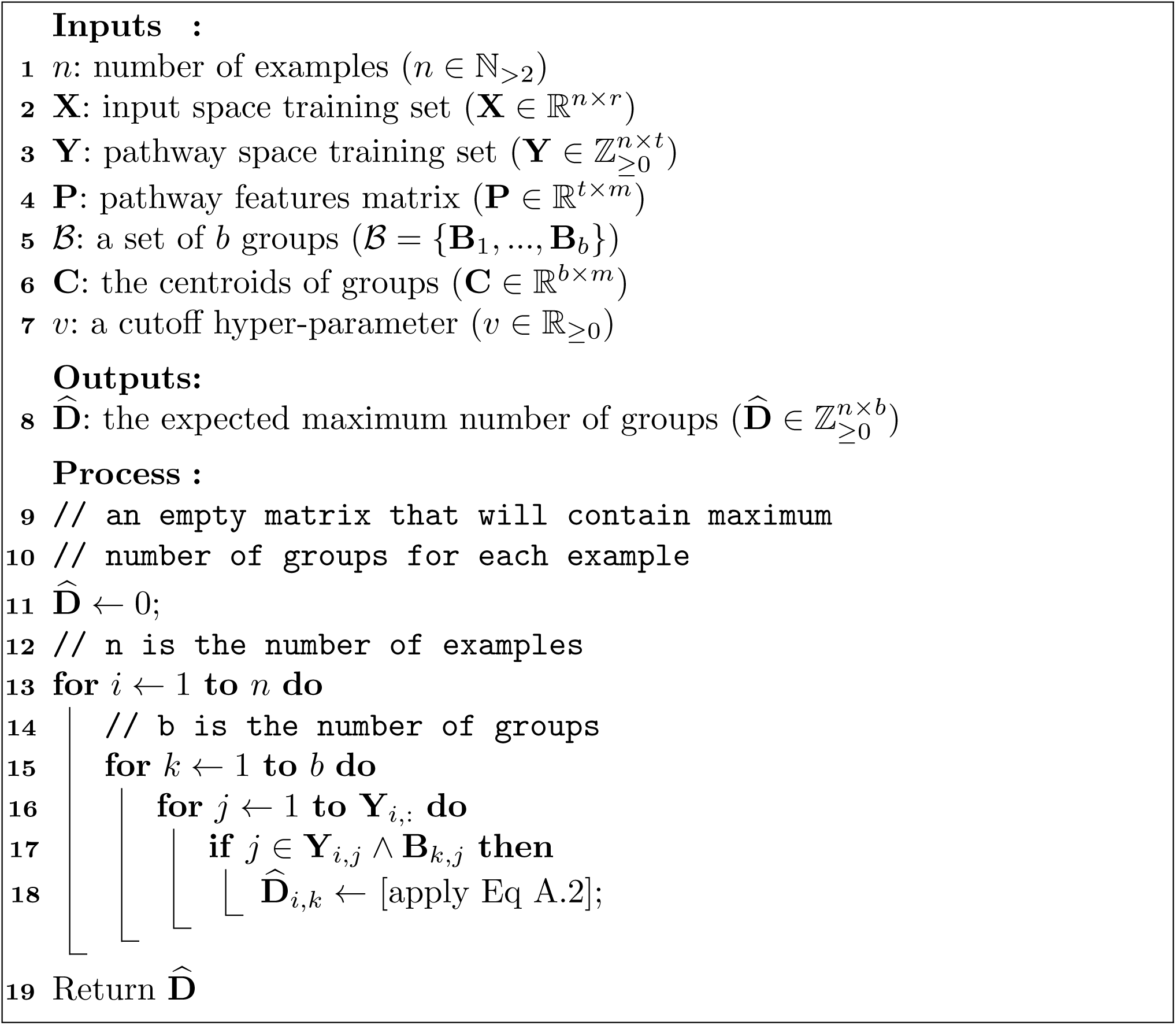

##### Constructing Pathway Group

In this step, pathways in *𝒮* are partitioned into non-disjoint *b* groups using any correlated models in Section B. These models are equipped to provide us with a group covariance matrix denoted by *Σ* ∈ ℝ^*b×b*^that is transformed to a correlation matrix *ρ* = *C*^−1^*ΣC*^−1^ where 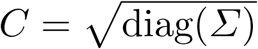 and a pathway distribution over groups denoted by *Φ* ∈ ℝ^*b×t*^. Each entry *Φ*_*i,j*_ corresponds to the probability of assigning a pathway *j* to the group *i*. For each group in *Φ*, we retain the top *k* (∈ 𝕫_≥1_) pathways based on the probability scores. The trimmed *Φ*^*!*^ ∈ ℝ^*b×k*^ (⊆ *Φ*) serves as an input to constructing centroids in the next step. Modeling pathway distribution and group correlation in this way are motivated by two key intuitions. First, organisms encoding similar pathways may share similar groups, thus, encouraging to have near-identical statistical properties for those organisms. Second, frequently occurring pathways in multiple organisms imply a similar relative contribution to a group.

##### Building Group Centroid

Having obtained a set of groups, reMap computes centroids for each group to capture the relative contribution of each pathway to its associated group’s centroid in the Euclidean space. Estimating centroids requires representing pathways and groups as vectors of real numbers. For this, we apply pathway2vec [10] to obtain pathway features. Then, the centroid of a group, say *s*, is computed according to:

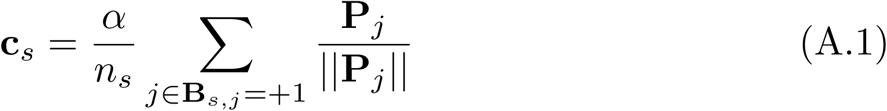

where **B**_*s*_ ∈ {−1, +1}^*t*^ is the group *s* obtained from the trimmed *Φ*_*s*_ after transforming it to {−1, +1}^*t*^. **c**_*s*_ corresponds the centroid of the group *s*, **P**∈ ℝ^*t×m*^ is a pathway representation matrix obtained from pathway2vec, *n*_*s*_ is the number of pathways (|{**B**_*s,j*_ = +1, ∀*j* ∈ *t*}|) in group *s*, ||.|| is the length of a feature vector, and *α* (∈ ℝ_*>*0_) is a hyper-parameter determined by empirical analysis (16 in this work). The proposed Eq. A.1 is based on the intuition that pathways associated with a group are semantically “close enough” to the center of the corresponding group, and the overlapping pathways among groups exhibit similar semantics with their associated groups. This procedure is described in Algorithm 1. In addition to the centroid computation, reMap also estimates a maximum number of expected groups to be annotated for a given example, indexed by *i*, using cosine similarity metric [9]:

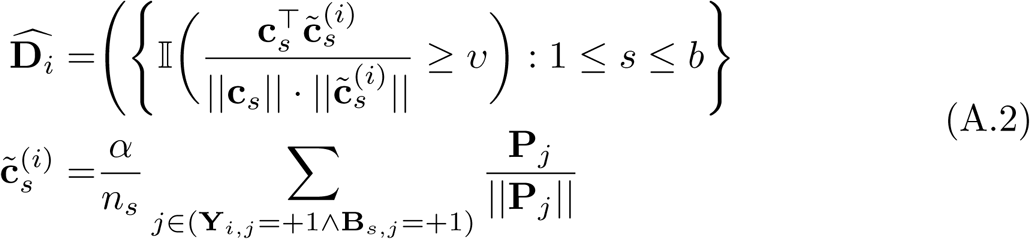

where 𝕀(.) is an indicator function that results in either +1 or −1 depending on a user-defined threshold *υ* (∈ ℝ_*>*0_). **Y**_*i*_ ∈ {−1, +1}^*t*^ corresponds to pathways either present or absent for the *i*th example, indicated by +1 and 1, respectively. 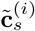 represents the centroid of the group *s* calculated based on pathways that are associated with the group *s* and are present in *i*th example. ñ_*s*_ is the number of pathways (|{**Y**_*i,j*_ = +1 ∧ **B**_*s,j*_ = +1, ∀*j* ∈ *t*}|) in group *s*. 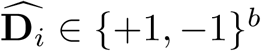 is a pre-optimized set of groups tagged for the *i*th example that will be used in the mapping step. Algorithm 2 describes the pseudocode for Eq. A.2.

##### Mapping Pathways to Pathway Groups

The goal of this step is to map an example to pathway groups, resulting in an optimized pathway group dataset 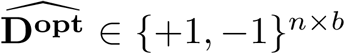. Formally, let us denote a set of groups that are picked to tag an example by 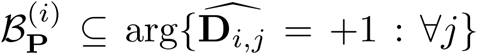 while the remaining unpicked groups is denoted by 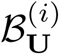, where 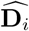 is obtained using Eq. 2.2. Both sets of groups are stored in 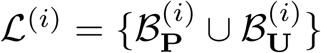. Then, reMap performs mapping in an iterative way, mirroring sequential learning and prediction strategy [18], where for each *i*th example, a group **B**_*j*_ at round *q* is either: i)-added to ℒ^(*i*)^, indicated by 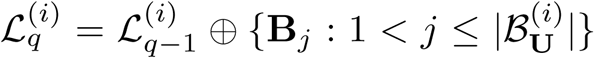; or ii)- removed from the set of selected groups, represented by 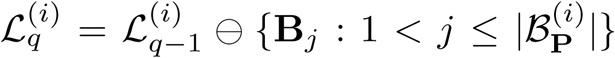. More specifically, at each iteration *q*, reMap estimates the probability of an example, given the selected groups that are obtained from the previous round *q* − 1, using threshold closeness (TC) metric [2] as:

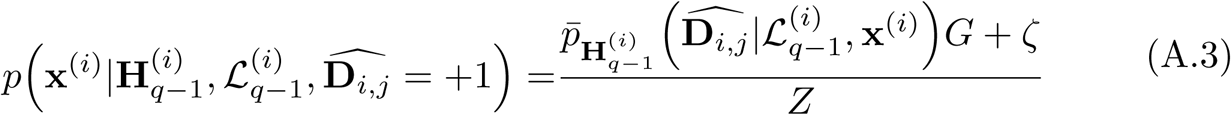

where **x**^(*i*)^ ∈ ℝ^*r*^and *r* is the total number of enzymes, 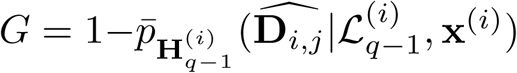 and 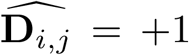 if the group **B**_*j*_ is tagged with the *i*th example. 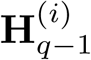 represents the history of prediction probability storing all 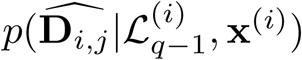 before the current iteration *q* while 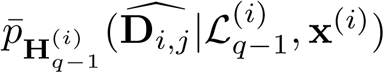 is the average probability of classifying **x**^(*i*)^ to the group **B**_*j*_ over values in 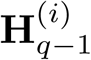. The term *ζ*(∈ ℝ_*>*0_) is a smoothness constant and *Z* is a normalization constant. Note that TC is a class conditional probability density function that encourages correct class probability to be close to the true unknown decision boundary. Hence, this step will ensure the correct latent group to be assigned to the *i*th example. To estimate 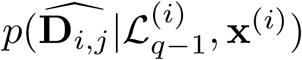, we jointly compute the probability of groups and pathways that are associated with 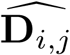 at round *q* − 1 as:

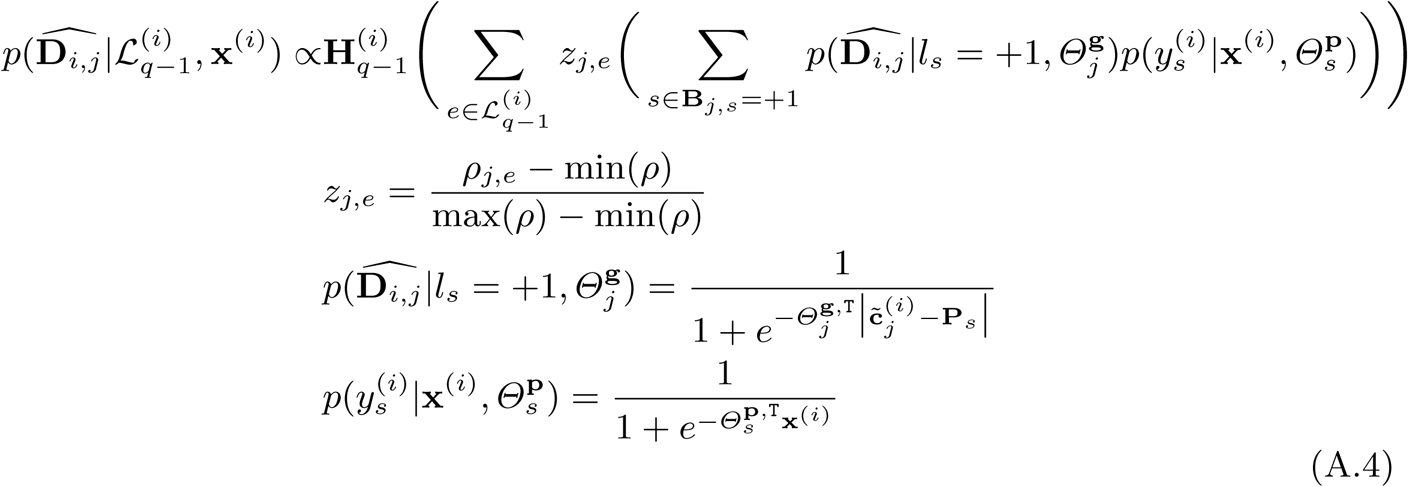

where 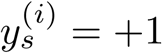 if the pathway index *s* is found to be present in both group *j* and in example **x**^(*i*)^ and 0 otherwise, and *l*_*s*_ = 1 if the pathway index *s* is associated with group *j* and 0 otherwise. *z*_*j,e*_ is a normalized correlation between groups *j* and *e*, respectively, obtained from *ρ* and 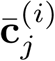 is presented in Eq A.2. 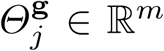 and 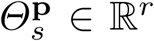 denote parameters for the group *j* and the pathway *s* model’s, respectively, and are learned during the feed-backward stage.

To reduce computational latency, instead of applying the above procedure to all groups for each example at every round, we randomly sub-sample groups of size *γ* (∈ 𝕫_*>*1_). Also, the estimate is still in the probability realm, therefore, we utilize a cut-off decision threshold (*β*) to retrieve a subset of groups having less overlapping pathways. Afterwards, ℒ^(*i*)^ will be updated either by adding or removing groups from a previous iteration. Algorithm 3 presents the pseudocode for relabeling multi-label dataset with groups.

###### Algorithm 3: Relabel2Group(*n*, **X, Y**,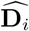,*𝔅*, **P**, *Θ*^**g**^, *Θ*^**p**^, **C**, *z, α, d*, ∈, *v, ζ, τ*)

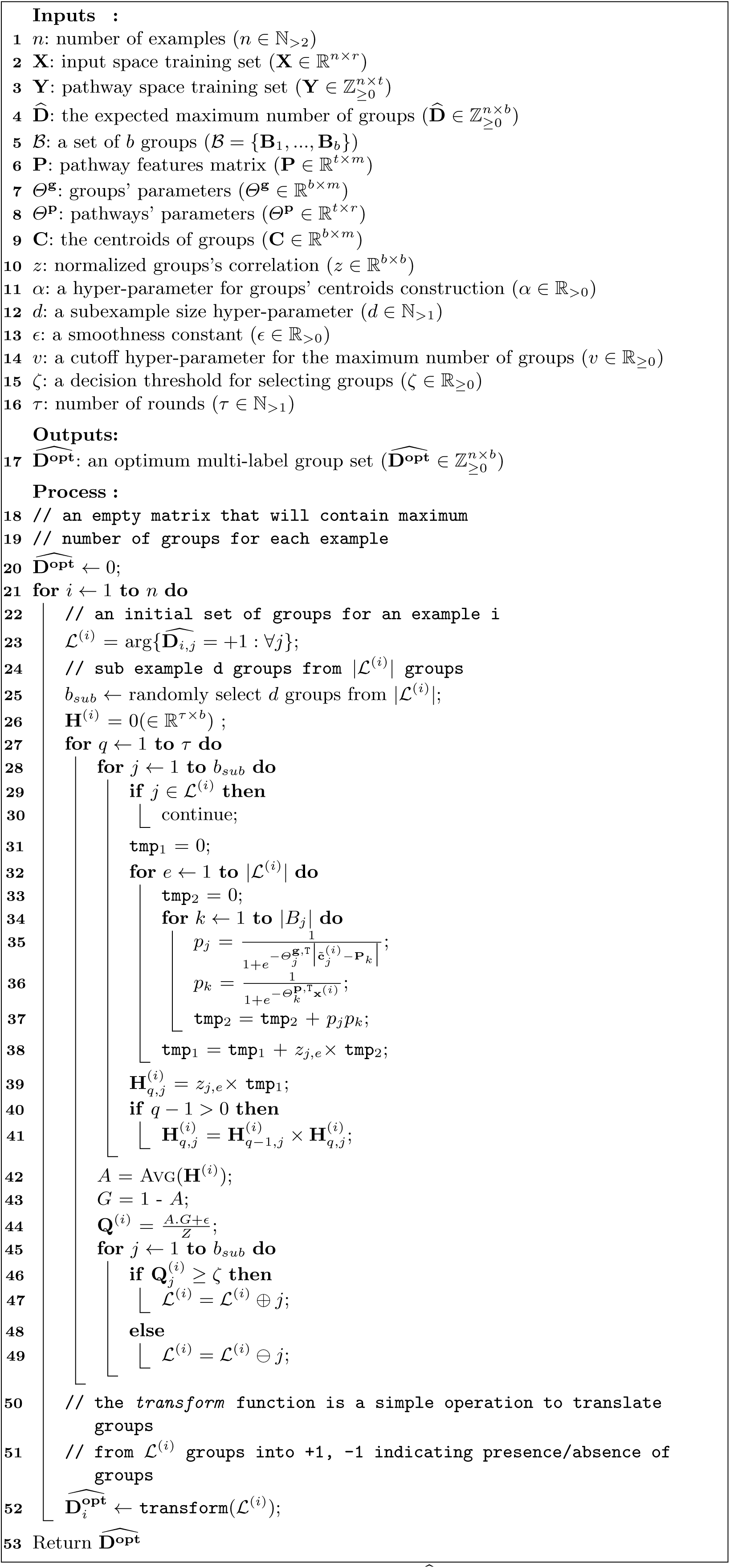

#### A.2 Feed-Backward Phase

Here, we set up the learning framework for computing reMap’s group and pathway parameters, jointly denoted as *Θ* = {*Θ*^**g**^, *Θ*^**p**^}. From Eq. A.3, three learning components can be identified: i)- a hyper-plane in the group space to absorb group correlation, ii)- a hyper-plane in the pathway space to encode semantic information about pathways, and iii)- a joint learning between groups and pathways to exploit pathway-group relationship. Let us define three empirical loss functions, corresponding the three components: ∈^**g**^ : {0, 1}^*b*^→ ℝ_≥0_, ∈ ^**p**^ : {0, 1}^*t*^ → ℝ_≥0_, and ∈ ^**gp**^ : {0, 1}^*b*^ → ℝ_≥0_ of margin **d***h*^**g**^(**x**), **y***h*^**p**^(**x**), and **d***h*^**gp**^(*y*), respectively, where *h*^(.)^ are decision functions. The last two loss functions are based on the logistic loss while the first loss is a sum of the two other losses. Now, to compute *Θ*, we maximize the posterior probability of Eq. A.4:

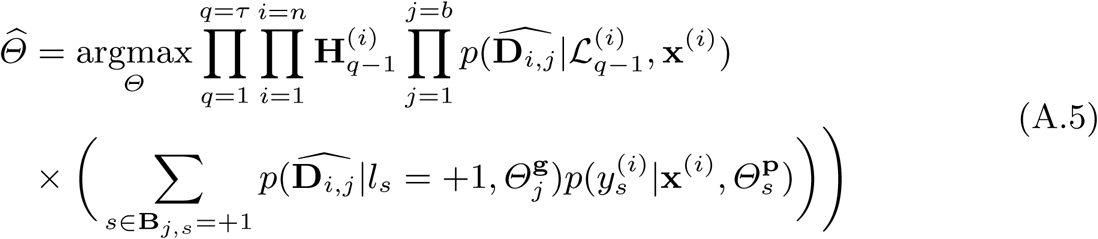

Estimation of parameters in Eq. A.5 is intractable due to the chain of probabilities 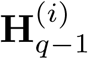 and the two marginalizations over ℒ_*q*−1_ and *s*. Hence, we propose the following two diagnoses: i)- conditional independence assumptions where the previous history values are independent given the most recent estimates and ii)- collapse the marginalization over ℒ_*q*−1_ by choosing only the maximum correlation *z*, irrelevant to which groups were considered. These simplified treatments provide an efficient way to optimize the parameters, where we adopt the “one-vs-all” scheme learning for each group and pathway [52].

In addition, we apply four constraints to retrieve a good set of parameters: i)- similarity between groups and associated pathways; ii)- weights of pathways, in a group, should be close to each other; iii)- examples sharing similar pathways should share similar representations; and iv)- all reMap’s parameters should not be too large or too small. These four constraints are important to allow smooth updates and mapping operations. Using these four constraints, the obtained pathway group dataset 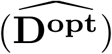, and the pathway data (**Y**), our objective function is formulated according to:

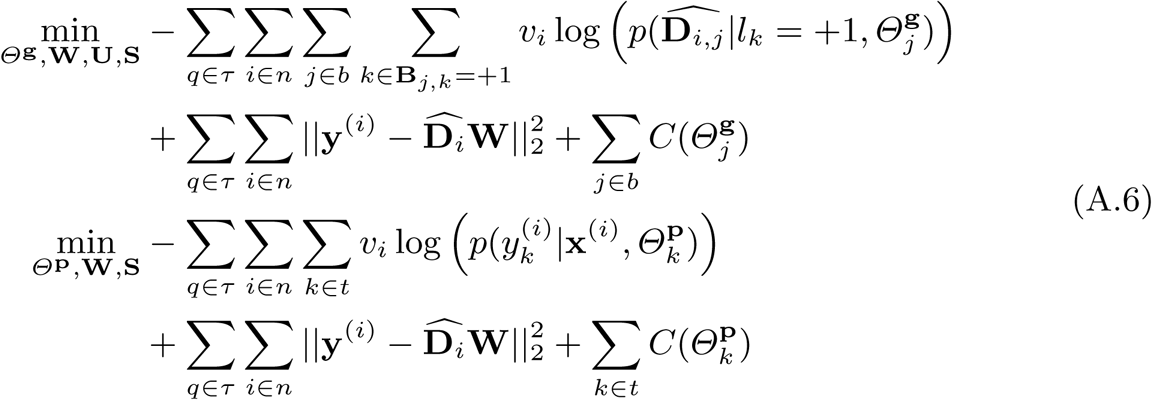

where

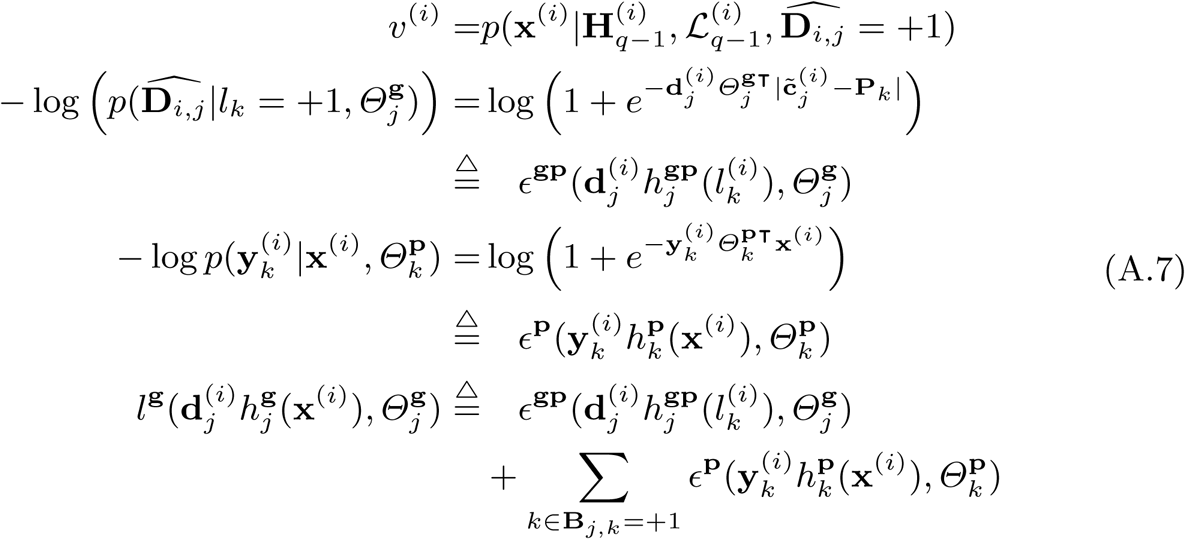

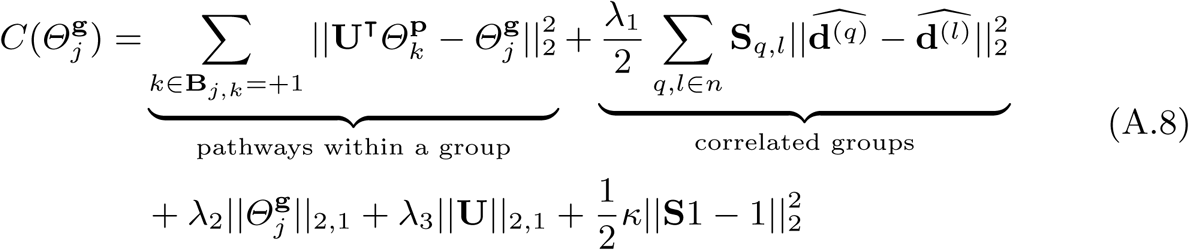

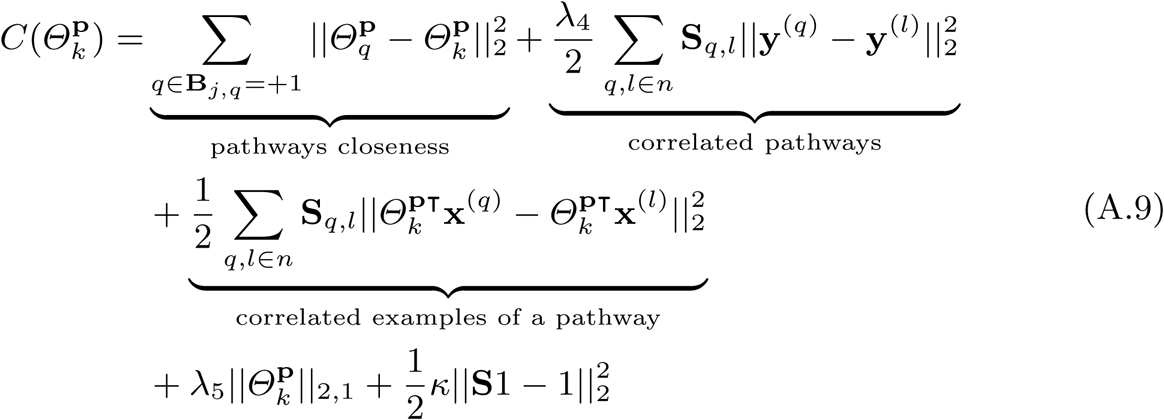

where 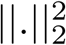 represents the squared *L*_2_ norm, 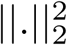 is the sum of the Euclidean norms of columns of a matrix, *v*^(*i*)^ is the weight of a example **x**^(*i*)^ to emphasize selection of informative examples, and *λ*_[1,2,3,4,5]_ ∈ ℝ are hyper-parameters controlling the relative contributions of the associated constraint terms. Let us explain all the terms involved in Eqs A.7-A.9. The function 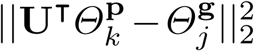 reflects the first constraint, where it enforces similarities between pathways, associated to a group *j*, and the pathway group *j* itself. **U** ∈ ℝ^*r×m*^ is the linear transformation matrix from *r* onto *m* dimensional space. For the second constraint, the term 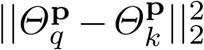 considers the similarities among pathways, grouped under a specific pathway group. To adopt the third constraint, we used four terms: 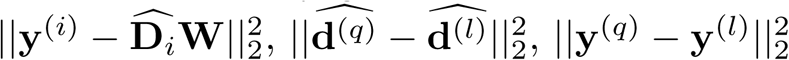 and 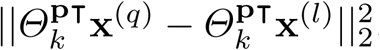.

The term 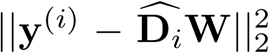 maintains the integrity of both pathway group and pathway vectors on example *i*, thus, encouraging groups to have similar contents as pathways, and **W** ∈ ℝ^*b×t*^ captures the correlation between groups and pathways. Both 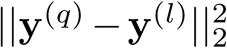 and 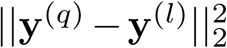 describes the resemblances between the two pathway group vectors, 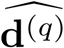 and 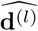, and the two pathway vectors, **y**^(*q*)^ and **y**^(*l*)^, suggesting the similarity between input instances **x**^(*q*)^ and **x**^(*k*)^. The similarity scores of the aforementioned instances are captured by 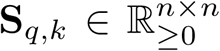, where a high score, indicates both examples have near identical pathways and, hence, should have similar groups, and vice-versa hold as well.

The formula 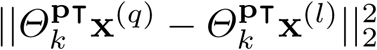 addresses the neighborhood relationship in the example feature space between **x**^(*q*)^ and **x**^(*k*)^, as characterized by **S**_*q,l*_ score [50]. As discussed before, if two instances are close to each other then they may possess relevant labels, which leads to relabeling a dataset with a proper subset of groups, hence, mitigating from the negative influences of imperfectly labeling groups. The terms 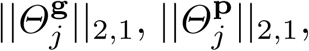, and ||**U**||_2,1_, constitute the fourth constraint that aim to shrink weights and perform feature selection. Finally, 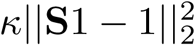 enforces equality constraint such that ∀*q*, ∑_*k*∈*n*_ **S**_*q,l*_ = 1, where *κ* is a Lagrange multiplier, and 1 denotes a column vector with all of it’s elements are equal to 1.

Taken together, the trainable parameters of reMap are: 1)- group-projection weight matrix **W**, 2)- pathway-projection weight matrix **U**, 3)- group-specific weight matrix *Θ*^**g**^, 4)- pathway-specific weight matrix *Θ*^**p**^, 5)- example-similarity specific weight matrix **S**, and 6)- group-specific updating matrix 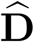. The last parameter is a binary matrix indicating the presence/absence of groups in the training dataset, which is gradually updated based on the gradient score strategy.

Unfortunately, the objective function in Eq. A.6 involves *L*_2,1_-norm that is non-smooth and difficult to be solved, instead we perform iterative gradient descent method for reMap which alternatively optimizes over one of six classes of variables (**W, U**, *Θ*^**g**^, *Θ*^**p**^, **S**, and 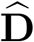) at a time while the others are held constant. The partial derivative of each term in Eq. A.6 is a positive semi-definitive, hence, the whole term is jointly convex, which leads to the following independent optimization problems for all pathways and groups classifiers according to the multi-label 1-vs-All approach [52].

**– Update W**. The gradient of Eq. A.6 w.r.t. **W** has the following formula:

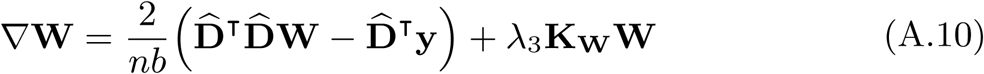

where 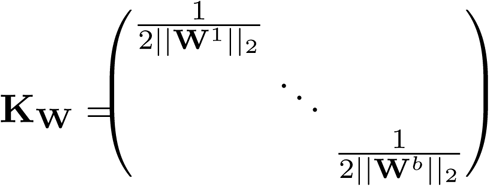
**– Update U**. The gradient of Eq. A.6 w.r.t. **U** becomes:

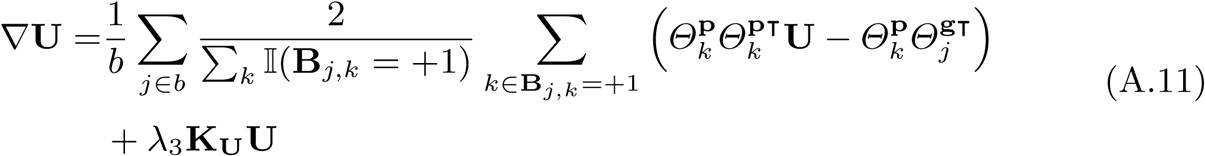

where 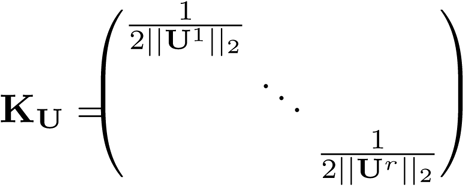
**– Update** *Θ*^**g**^. The partial derivative for each pathway group, say 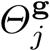, is:

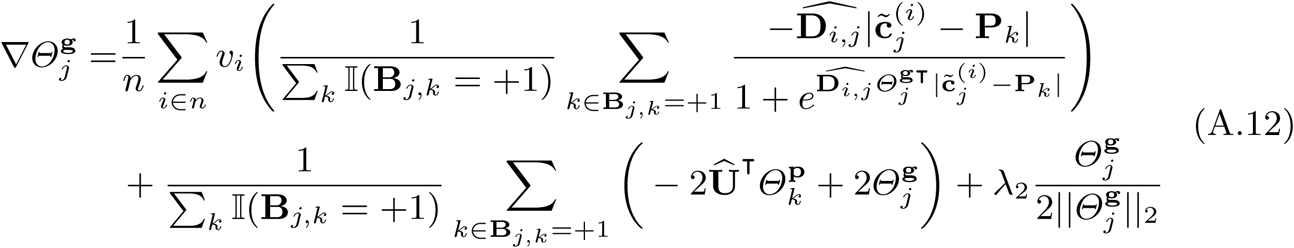

where Û obtained from Eq. A.11.
**– Update** *Θ*^**p**^. The partial derivative w.r.t one pathway *k* of *Θ*^**p**^ with the new Û and 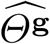 updates has the following form:

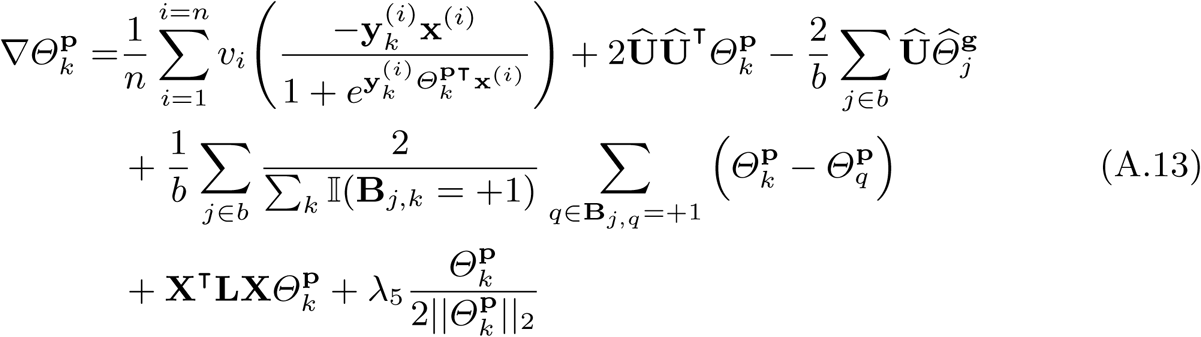

where **L** ≜ **M** − **S** is the graph Laplacian matrix and **M** is a diagonal matrix with **M**_*j,j*_ = ∑ **S**_*j,k*_. Note that 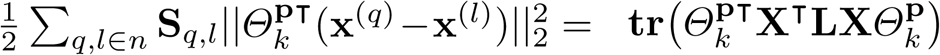. Following the work of [44], it is important for practical purpose to normalize the graph Laplacian, to account for the fact that some examples are more similar than others [48]: 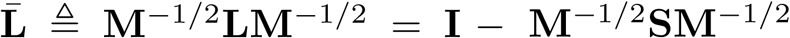. Adhering to this property, we consider the following formula: 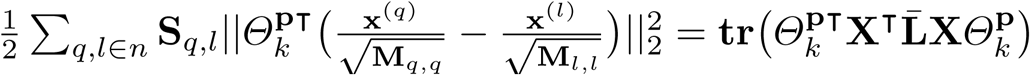
**– Update S**. Given the updated values of Û and 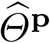, we obtain the equivalent objective function of Eq. A.6 with the terms only related to **S** as:

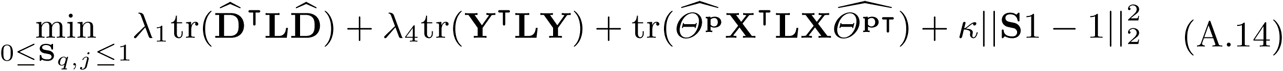 For the inequality constraint, during iterative updates we force values of **S** to be within the range of [0, 1]. Then, the gradient update can be written as:

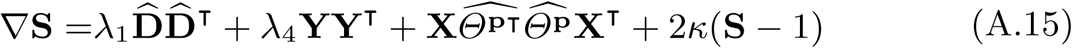

As we have mentioned, the similarity matrix **S** captures reliable and discriminative locality information in the projected example feature space, and this information is utilized to optimize correlations in the predicted pathway space, which ensures example-pathway space consistency. Consequently, a set of groups can be inferred with high fidelity, for each example, if features with labels correlation information is disseminated to features with groups correlations, thus, alleviating the effects of imperfectly detecting negative groups.
**– Update** 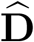. We iteratively update groups, where a set of groups are added or removed at each round. In particular, a positive subset of groups is selected 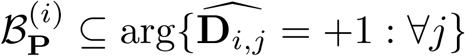, for each example **x**^(*i*)^, and the remaining groups are considered to be negative to that example. While it is relatively easy to compile a set of positive groups for an example, however, groups not belonging to that example are too diverse to be considered as negative. Thus, it is better to consider the remaining groups as unassigned 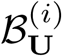. We use the gradient score strategy, where the values of 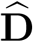 is updated based on the gradient score according to:

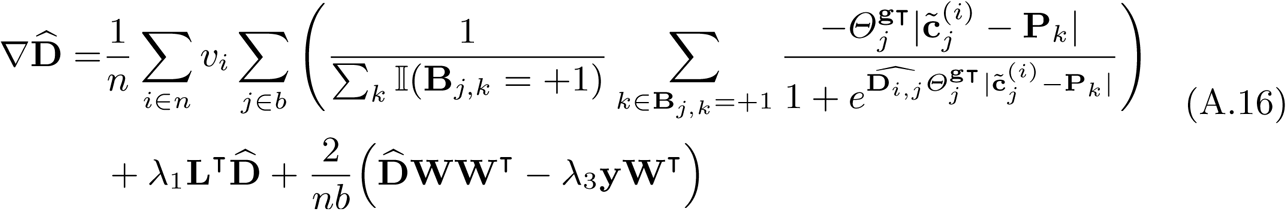

After getting the gradient score, we assign groups to examples based on:

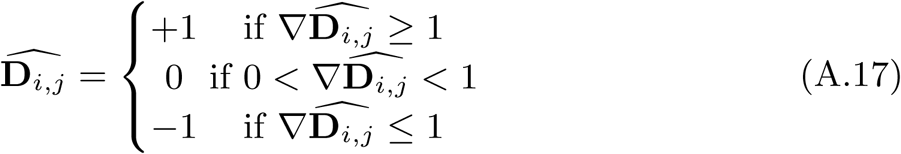

where 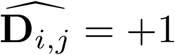 (resp. −1 and 0) means the group is selected to be positive (resp. negative and unknown) given a training example *i*. Having acquired a new selected set of groups for each instance, we update ℒ^(*i*)^ accordingly. The pseudocode for this phase in presented in Algorithm 4

#### A.3 Closing the loop

The two phases are repeated for all examples in a given pathway data, until a predefined number of rounds *τ* (∈ 𝕫_*>*1_) is reached. At the end, a pathway group dataset is produced which consists of *n* examples with the assigned groups, i.e., 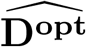. This data can be used as inputs to a pathway predictor to perform pathway prediction for a newly sequenced genome.

##### Algorithm 4: Backward(*n*, **X, Y**, 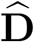, **P, C**, *z, d, ξ, λ, γ*)

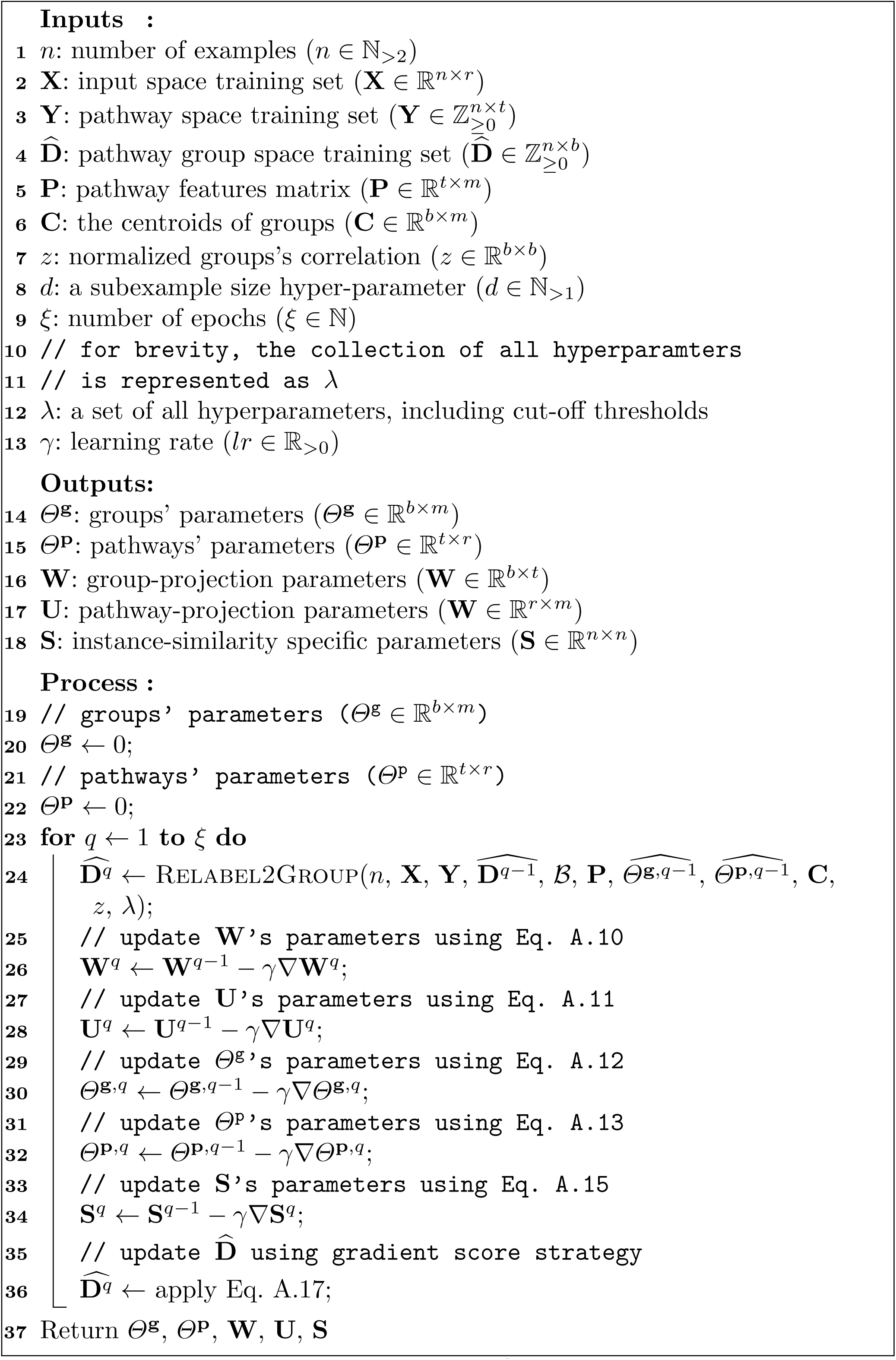

### B Correlated Models

We present three correlated pathway models that can be applied during pathway group construction step in the feed-forward phase of reMap: i)-CTM (<u>c</u>orrelated <u>t</u>opic <u>m</u>odel) [23], ii)- SOAP (<u>s</u>parse <u>c</u>orrelated <u>p</u>athway group) and iii)- SPREAT (di<u>s</u>tributed s<u>p</u>arse cor<u>re</u>lated p<u>a</u>thway group). These models incorporate pathway abundance information to encode each example as a mixture distribution of groups, and each pathway group, in turn, is a mixture of pathways with different mixing proportions. The pathway abundance information can be obtained by mapping enzyme –with abundances– onto the reference pathway database (e.g. MetaCyc). Before we discuss these three models, first let us provide some background information and notations. We note that each mathematical symbol is only related in the context of this section.

#### Definition 6. Pathway Collection

*Let* 𝒫 = {**y**^(*i*)^ : 1 *< i* ⩽ *n*} *be a collection of n examples, where each example* 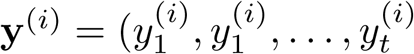 *is a vector encoding the unnormalized abundance information of pathways and t is the pathway size. Let 𝒴* = {*h*_1_, *h*_2_, …, *h*_*t*_} *be a set of all known metabolic pathways obtained from a trusted source (e*.*g*., *MetaCyc [28]), and 𝒴* ⊆ *𝒴 corresponds to a subset of true pathways associated with the example i*.

Recovering latent distributions of *𝒫* mirrors the concept modeling paradigm, which aims to reconstruct the thematic structure, called “topics”, from a corpus [25].

#### Definition 7. Concept Modeling

*Given a collection of n examples, a concept distribution for i-th example is a multinomial distribution vector, denoted by η*^(*i*)^ *of size b concepts, i*.*e*.,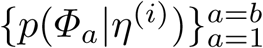, *where Φ*_*j*_ *in a multinomial feature distribution over the concept j, i*.*e*., 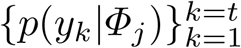. *The overall goal of concept modeling is to recover the b salient concepts of each example*.

In this paper, the term concept is referred to as “pathway group” or “group”. For brevity purposes, the following terms: *concept, topic*, or *pathway group*, are used interchangeably. Also, *features* correspond to *pathways*.

The classical studies in concept modeling attempt to discover concepts from a collection of examples that are composed of features, as in the case of latent Dirichlet allocation (LDA) [25]. However, this approach neglects dependencies among concepts. We take advantage of the inherent thematic structure of examples and model the concept dependencies to extract the concept distributions of examples.

#### Definition 8. Concept Correlation

*Given 𝒫, the pairwise concept-correlation is defined by a Gaussian covariance matrix, denoted by Σ. Each entry s*_*i,j*_ *in Σ characterizes the i-th pathway group association with the pathway group j, where a larger score indicates both concepts are highly correlated*.

However, there exist situations where a set of pathways may not be included in *𝒫* because *𝒫* has high noise. An alternative way to incorporate missing pathways is to store these pathway in a separate list while keeping the original pathway collection intact for further investigation. Lets us denote 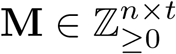 a matrix holding a set of missing pathways where each entry is an integer value indicating the abundance of a pathway in an example. Here, this matrix is referred to as a “background” or “supplementary” matrix, analogous to studies in [32, 53]. With these definitions, we describe the correlated models.

#### B.1 Correlated Topic Model

The <u>c</u>orrelated <u>t</u>opic <u>m</u>odel (CTM) is a probabilistic graphical model that extends the generative story of LDA [25] to incorporate correlation among concepts. Fig. 6a shows the Bayesian graphical model for CTM using plate notation. Like latent Dirichlet allocation [25], the CTM is comprised of a hierarchical Bayesian mixture model, where features (words as described in the original paper) are mixed to constitute concepts. And, the concepts are assumed to be correlated to each other by a Gaussian covariance matrix.

**Fig. 6:**
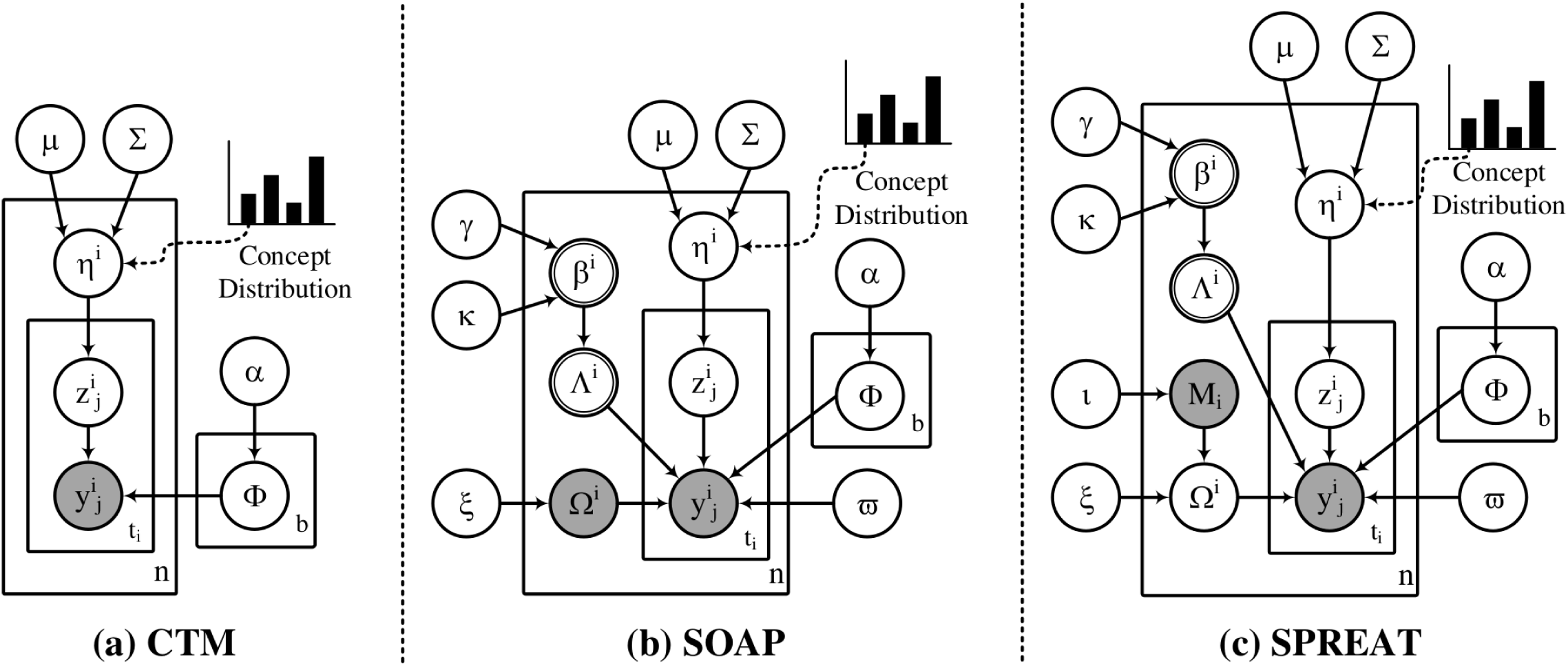
Graphical model representation of the correlated concept models. The boxes are “plates” representing replicates. The outer plate represents instances, while the inner plate represents the repeated choice of features within an example. The logistic normal distribution, used to model the latent concept proportions of an example, captures correlations among concepts that are impossible to capture using a single Dirichlet. The observed data for each example **x**_*i*_ are a set of annotated features **y**_*i*_ and a set of hypothetical features **M**_*i*_. The hidden variables are: per-example concept proportions *η*_*i*_, per-example concept selection parameters *Λ*_*i*_, per-example hypothetical feature distributions *Ω*_*i*_, perfeature concept assignment *z*_*i,j*_, per-concept distribution over features *Φ*_*a*_, and per-feature indicator parameter *d*_*i,j*_.

Formally, let *n* be the total number of a collection, where each example *i* consists of features, i.e., **y**^(*i*)^. Then, the generative process for CTM is described as follows. First, we draw a multinomial feature distribution *Φ*_*a*_ from a Dirichlet prior *α >* ℝ_*>*0_ for each concept *a* ∈ {1, …, *b*}. Then, for each example *i*, a Gaussian random variable is drawn *η*^(*i*)^ ∼ *N*(*μ, Σ*), where *μ* is a *b* dimensional mean and *Σ* ∈ ℝ^*b×b*^is the covariance matrix. The random variable *η*^(*i*)^ is projected onto the probability simplex to obtain the concept distributions *θ*^(*i*)^ = softmax(*η*^(*i*)^), corresponding the logistic-normal distribution, from which a concept indicator 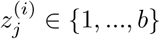 is sampled. Finally, each observed feature *j* ∈ {1, …, *t*^*i*^} is drawn from the associated feature distribution, indicated by it’s concept assignment, i.e.,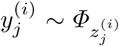. This generative process is outlined in Algorithm 5, which can be observed that the process is identical to LDA except the concept distributions is sampled from the logistic normal rather than a Dirichlet prior.

##### Algorithm 5: The generative process for CTM given a collection

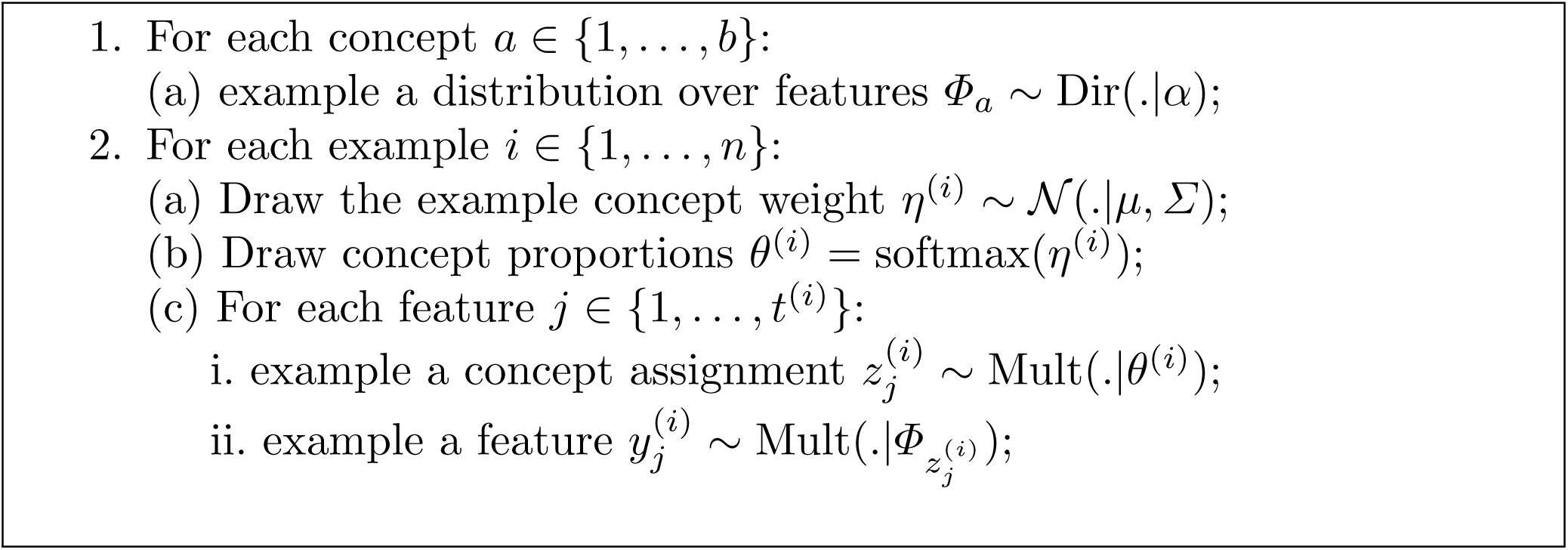

#### B.2 Correlated Pathway-Group Model

Correlated pathway group models are extension to CTM (Figs 6b and c): i)SOAP and ii)- SPREAT. Both models incorporate dual sparseness and supplementary pathways in modeling group proportions. These important properties are not adopted in CTM. Let us discuss these two models.

Analogous to CTM, given *n* number of examples and a matrix encoding the missing features **M**, the generative process for SOAP and SPREAT can be described as follows. First, we draw a multinomial feature distribution *Φ*_*a*_ from asymmetric Dirichlet prior *α* ∈ ℝ_*>*0_ for each concept *a* ∈ {1, …, *b*}, where *b* is assumed to be known and fixed in advance. The symmetric assumption is appropriate, in such a scenario, because our prior knowledge, associated with these features, is inaccessible. For each example *i*, a concept proportion is drawn *θ*^(*i*)^ = softmax(*η*^(*i*)^), where *η*^(*i*)^ is a Gaussian random variable with mean and covariance are denoted by *μ* and *Σ*, respectability.

To sample a concept, it is reasonable to expect that each example is usually explained with a handful set of a mixed proportion of concepts. Besides, a concept should cover a few focused features, instead of absorbing all features. Thus, we borrow the idea from [21,22,30,36,43] to enforce dual sparsity to retain those relevant focused concepts and features by: i)- introducing an auxiliary Bernoulli variable *Λ*^(*i*)^ of size *b* to determine whether a concept is selected for an example *i* or ignored, and ii)- applying a cutoff threshold to retain top *k* « *t* features for each concept. Instead of sampling each entry in *Λ*^(*i*)^ directly from a Bernoulli coin toss, we assume that each entry is sampled from a Beta distribution *β*^(*i*)^, parameterized by two hyperparameters *γ* ∈ ℝ_*>*0_ and *κ* ∈ ℝ_*>*0_. Applying this dual sparsity, we aim to enhance the interpretability of the learned concepts while reducing the negative correlation among concepts on *Σ*.

Next, a concept indicator 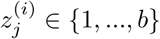 is drawn according to the example-specific mixture proportion *Λ*^(*i*)^ ⊙ *θ*^(*i*)^, where ⊙ represents the Hadamard product. Now each feature 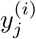 in example *i* is generated from a weighted distribution 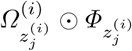, as indicated by it’s concept assignment, using a smoothing prior *ϖ* ∈ ℝ_*>*0_. The parameter *Ω*^(*i*)^ ∈ ℝ^*t*^, derived from **M**_*i*_, represents a normalized supplementary feature of size *t*, which is assumed to be drawn from a symmetric Dirichlet prior *ξ* ∈ ℝ_*>*0_. For SPREAT, this parameter encodes distribution, where each element of 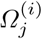 corresponds to the example’s probability of using feature *y*_*j*_ ∈ **M**_*i*_. Here, the background feature is assumed to be drawn from a sparse binary vector prior *ι* ∈ ℝ_*>*0_ that is included for completeness because each example’s feature **M**_*i*_ is already observed.

##### Algorithm 6: The generative process for SOAP and SPREAT

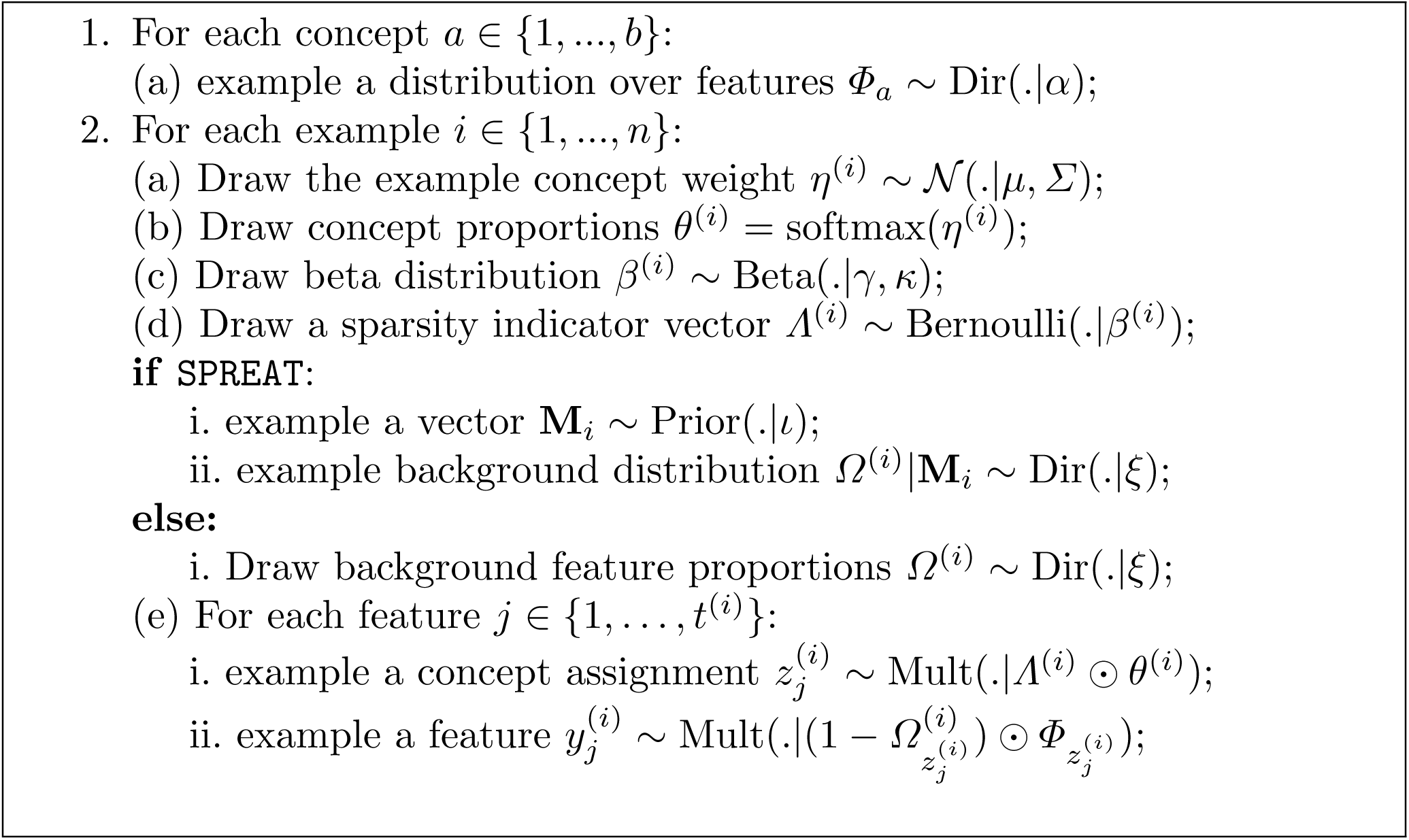

Representing SOAP and SPREAT as layer-wise mixing components supports the hierarchical modularity of metabolic pathway generation, where the components of one level (e.g., features) permit to contribute to other structures with different degrees of granularity. The generative process of SOAP and SPREAT models is summarized in Algorithm 6. Note that by setting all entries in *Ω, Λ*, and to 1, SOAP and SPREAT are reduced to CTM (“collapse2ctm” or c2m), which is an additional benefit to the these models.

#### B.3 Evidence Lower Bound (ELBO) for SPREAT

Here, we discuss the inference for the SPREAT model. Similar expression is straightforward to derive for SOAP. Given *𝒫*, the goal of inference is to compute the posterior distribution of the per-example concept proportions *η*^(*i*)^, the per-example concept selection parameters *Λ*^(*i*)^ and the associated beta distributions *β*^(*i*)^, the per-example background feature distributions *Ω*^(*i*)^, the per-feature concept assignment 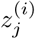, and the per-concept distribution over features *Φ*_*a*_.

Looking at the topology of the Bayesian network, we can specify the complete-data likelihood, i.e., the joint distribution of all observed and latent variables given the hyperparameters and sparse supplementary feature matrix following the model’s independence assumptions:

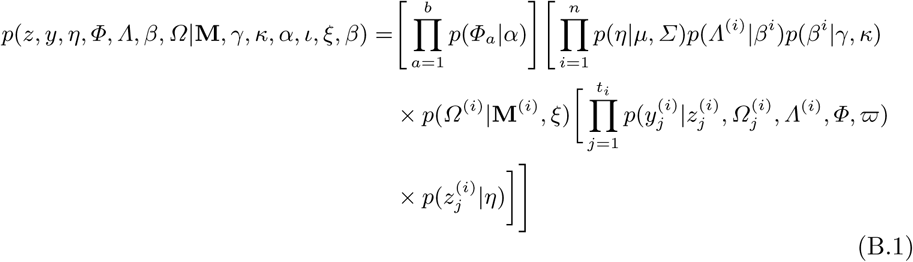

By denoting all parameters as *Θ* and variables as **V** while omitting hyper- parameters, we obtain the following posterior expression:

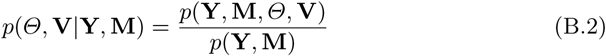

Unfortunately, the exact posterior distribution of the latent variables is computationally intractable. The numerator is easy to compute for any configuration of the hidden variables and parameters. The problem is the denominator, which is the marginal probability of the data:

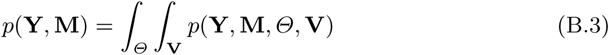

Computing the marginal requires a complicated integral over *n* examples of |*Θ*| parameters and another integral over the |**V**|^*n*^ configurations multiplied by the size of each variable in **V** As such, we appeal to the variational inference algorithm [25]. The main intuition behind variational methods is to first posit a family of distributions over the hidden parameters and variables that are indexed by a set of free parameters, and then fitting the parameters to find the member of the family that is closest to the true posterior of interest in Eq. B.2. The closeness is commonly measured using Kullback–Leibler (KL) divergence [35]. The resulting variational distribution is simpler than the true posterior so that the solution can be approximated. However, directly minimizing the KL divergence is infeasible due to the same reason that the posterior is difficult to compute, but, we can optimize an objective function that is equal to the negative KL divergence up to a constant. This is known as the evidence lower bound (ELBO), a lower bound on the logarithm of the marginal probability in Eq. B.3, i.e., log *p*(**Y, M**). This ELBO can be defined using Jensen’s inequality on a variational distribution over the hidden variables *q*(*Θ*, **V**) as:

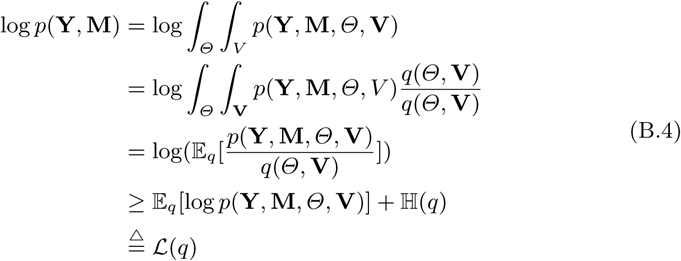

The ELBO contains two terms. The first term, 𝔼_*q*_[log *p*(**Y, M**, *Θ*, **V**)], captures how well *q*(*Θ*, **V**) describes a distribution of the model. The second term is the entropy of the variational distribution, 𝔼_*q*_[*−*log *q*(*Θ*, **V**)], which protects the variational distribution from “overfitting” [24]. Both of these terms depend on *q*(*Θ*, **V**), the variational distribution of the hidden variables.

The simplest variational family of distributions is the mean-field family where each hidden variable/parameter is fully-factorized and governed by its own parameter. This allows us to tractably optimize the parameters to find a local minimum of the KL divergence. For SPREAT, the mean-field variational distribution is expressed as:

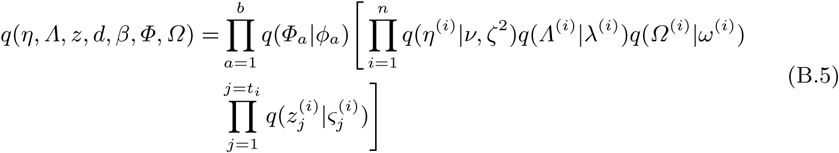

where *Φ, ν, ζ*^2^, *λ, ω* and *ς* are variational free parameters. Table 3 shows the correspondence between variational and the original parameters.

**Table 3:**
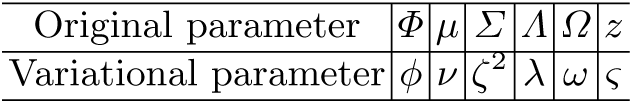
Correspondence between variational and original parameters.

Taking together, the first term in Eq. B.4, 𝔼_*q*_[log *p*(**Y, M**, *Θ, V*)], can be decomposed into:

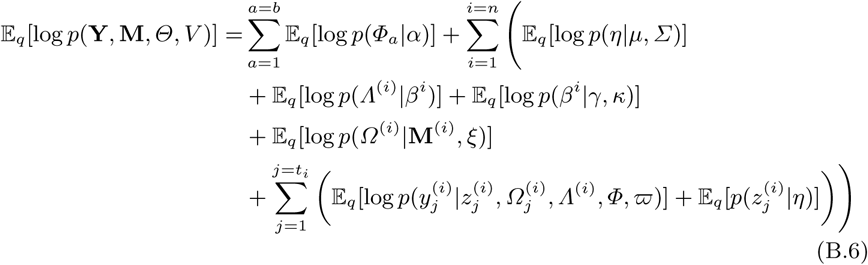

The second term ℍ(*q*) in Eq. B.4 can be expressed as:

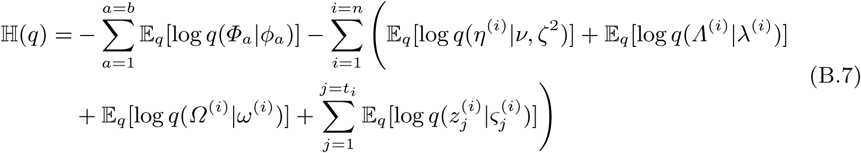

##### Variational Lower Bound

Given Eq. B.6, we derive expressions for each term:

1. For the concept distribution over features, which are Dirichlet-distributed,

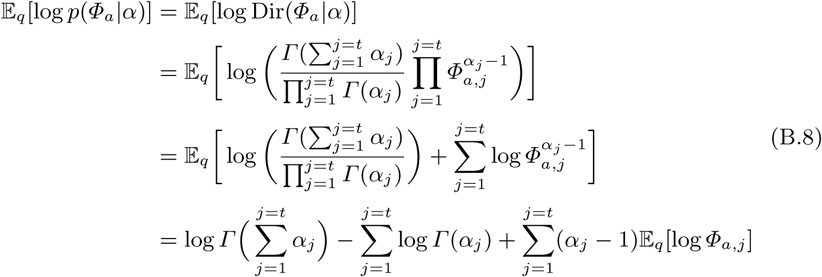
2. For the concepts probabilities for each example, which are Gaussian distributed,

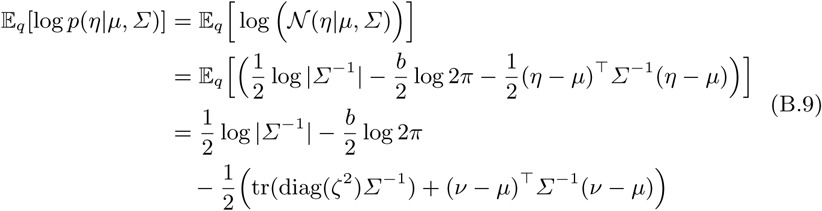
3. For the focused concept distributions for each example, which are Bernoulli distributed,

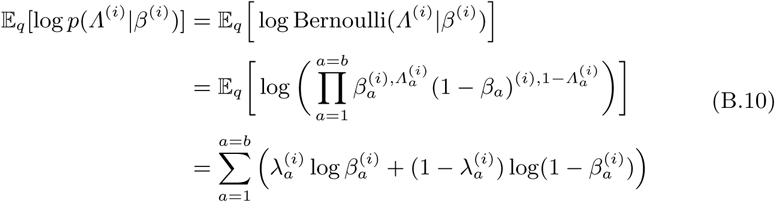
4. For selecting a set of focused concepts for each example, which are beta distributed,

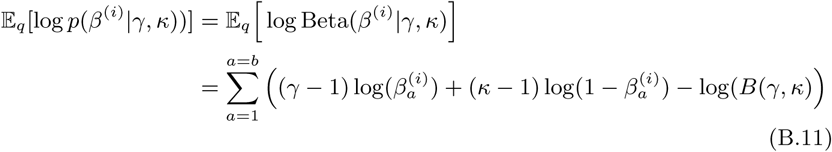
5. For the hypothetical feature distributions for each example, which are Dirichlet distributed,

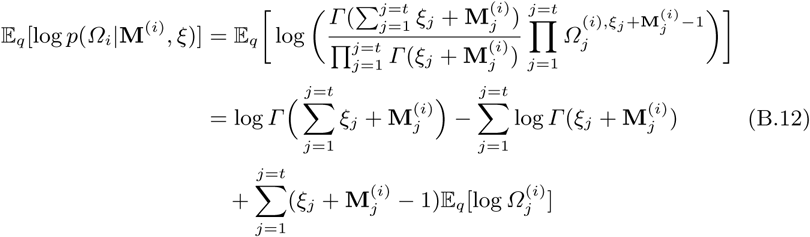
6. For the feature assignments from both concept-feature and hypothetical feature distributions,

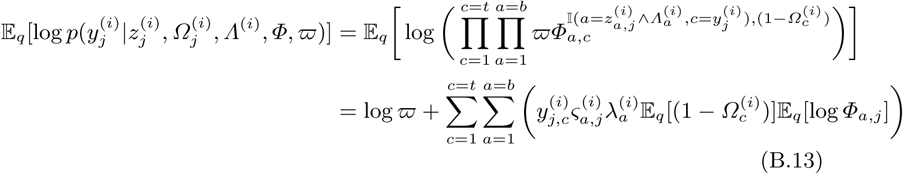
7. For the concept assignments over features, the expectation of the log probability of the latent concepts is given by:

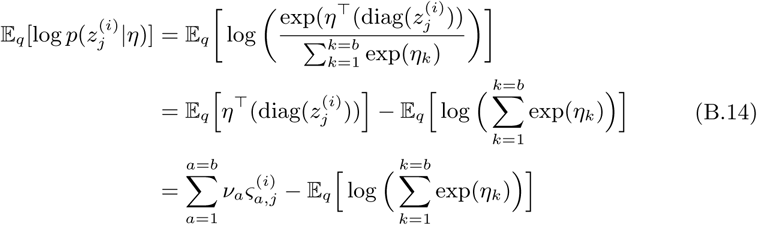

The second term is hard to compute, hence, we use the solution suggested by [23] in order to obtain the tightest lower bound on 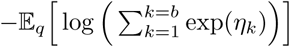 using a first-order Taylor expansion. Because the function *−*log is convex, a first-order Taylor expansion about the point *(2*, a variational parameter, produces the following inequality:

Plugging back the results into Eq. B.14, we obtain:

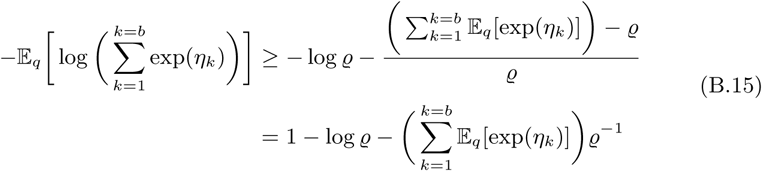

Now, for the entropy ℍ(*q*) in Eq. B.7, we decompose their expectations as:

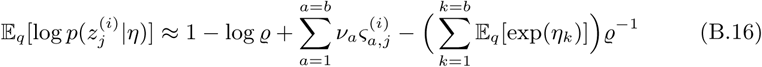

1. For the concept-feature distributions, which are Dirichlet distributed,

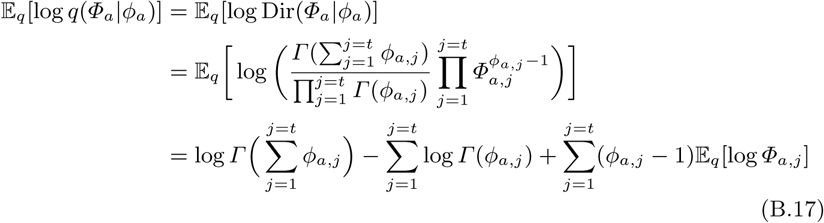
2. For the concept distributions, which are Gaussian distributed,

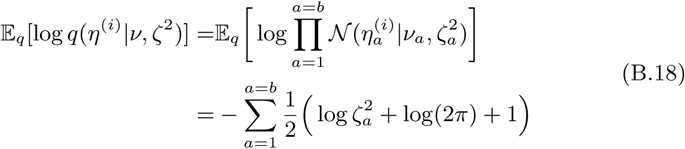
3. For the concept choice parameter, which are Bernoulli distributed,

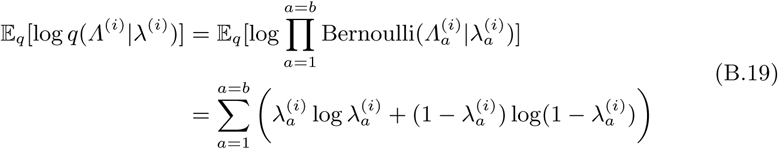
4. For the supplementary feature distributions over examples, which are Dirichlet distributed,

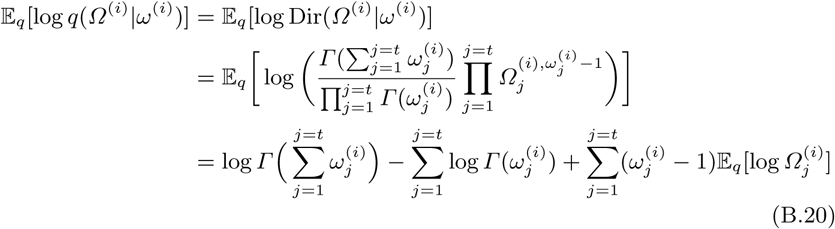
5. For the feature assignments over examples, which are multinomially distributed,

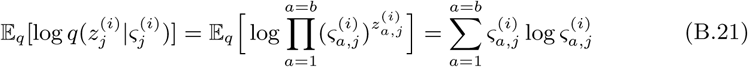

where the exceptions of all the above equations can be derived using:

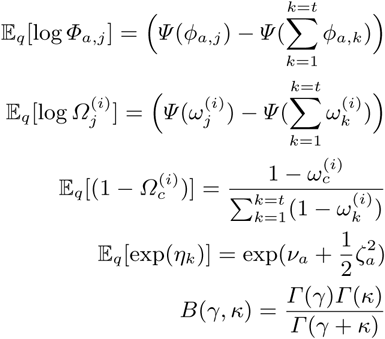

Not that *Γ* denotes the Gamma function while *Ψ* is the logarithmic derivative of the Gamma function.

##### Merging All the Expectations of the ELBO Terms

Now, by joining all the terms, the full ELBO can be defined as:

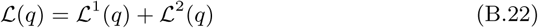

where,

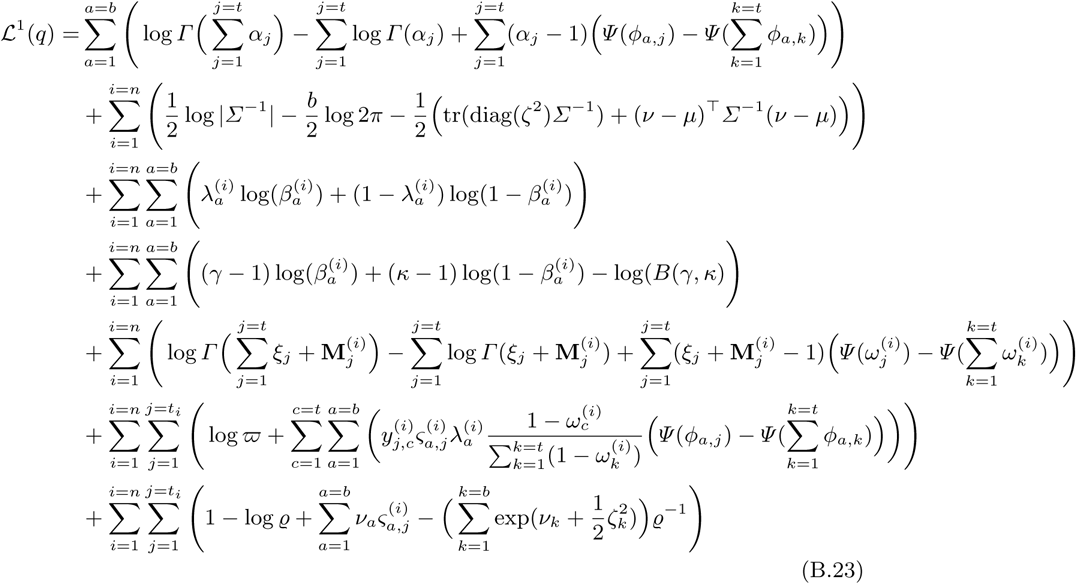

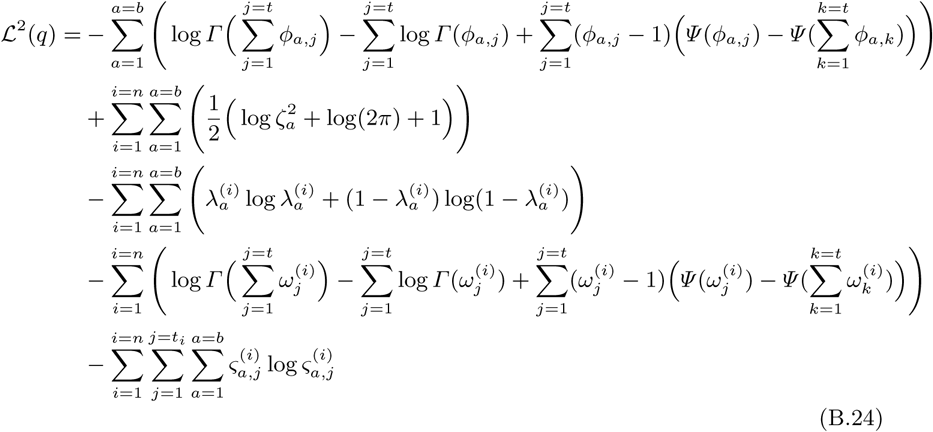

#### B.4 Optimizing the ELBO Terms

In this section, we maximize the bound in Eq. B.22 with respect to each variational parameters using coordinate ascent updates. Using this approach, each variational parameter is optimized individually while holding the remaining variables fixed. Practically, a more convenient way is to apply the mini-batch gradient approach that alternates between subsampling a batch of examples and updating each variational parameter, after being scaled by a learning rate [31]. This structure of learning assists us to approximate the posterior with massive examples, making the complete problem computationally scalable.

##### 1. Optimizing w.r.t. *ς*. Gathering only the terms in the bound that contain *ς*, we obtain

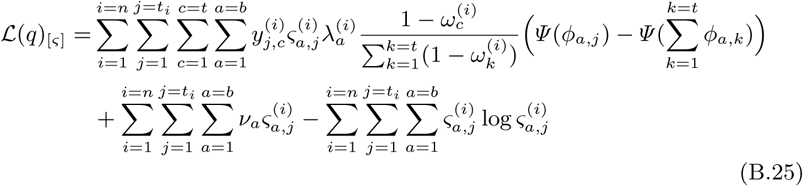

Taking derivatives w.r.t. 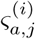, we obtain:

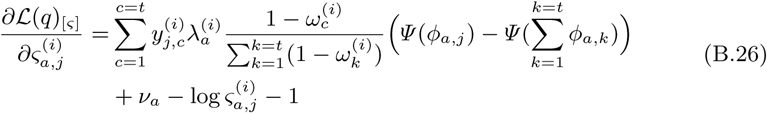

The analytical expression of the variational concept assignment *q*(*ς*) for each feature *yj* and concept *a* is not amenable due to the non-conjugacy of logistic-normal with latent variables. Instead, we approximate the solution according to:

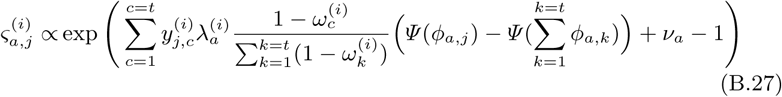

where *Ψ* (.) is the digamma function. Observe how the variational parameter 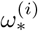 serves as the smoothing term in selecting concepts for each feature, either from **M***i* or from *𝒫*, when 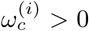. However, if 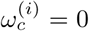, then 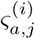 is updated based on *ϕa,j*.

##### 2. Optimizing w.r.t

*v*. Collecting only the terms in the bound that contain *v* gives,

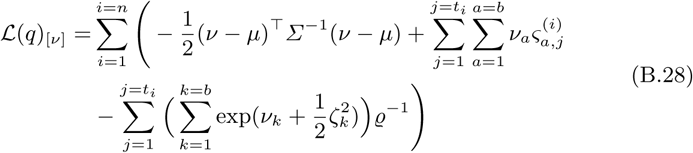

Taking derivatives w.r.t. *va* for each concept *a*, we obtain:

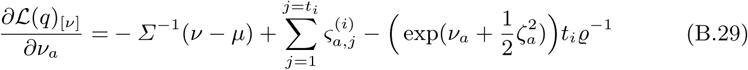

where *ϱ* is another variational parameter, as in CTM [23]. However, the above equation in hard to optimize, instead, we use a conjugate gradient algorithm.

##### 3. Optimizing w.r.t

ζ2. By symmetry, we gather all the terms that has ζ2 from Eq. B.22:

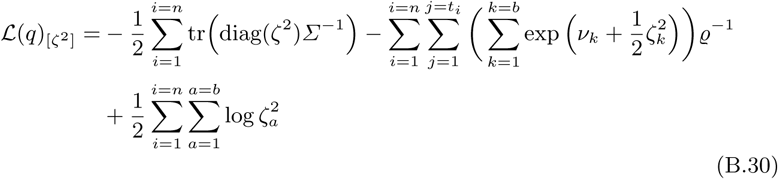

Taking derivatives w.r.t. 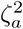 for each concept *a*, we obtain:

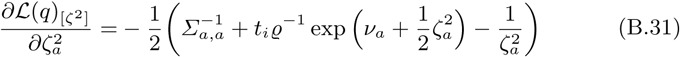

Again, there is no analytical solution to the above formula. Instead, we use the Newton’s method for each coordinate such that ζ*a* ∈ ℝ*>*0.

##### 4. Optimizing w.r.t

*ϱ*. Extracting the terms involving *ϱ* in the bound gives,\

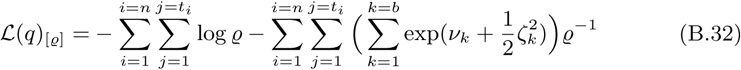

**Basher** et al.

Taking derivatives w.r.t. *ϱ*, we obtain:

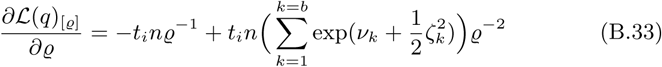

Equating the above formula to zero to obtain a maximum, we get:

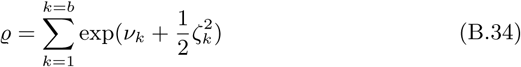

##### 5. Optimizing w.r.t

*ω*. Isolating only the terms in the bound that contain variational background feature distributions *q*(*ω*), we obtain:

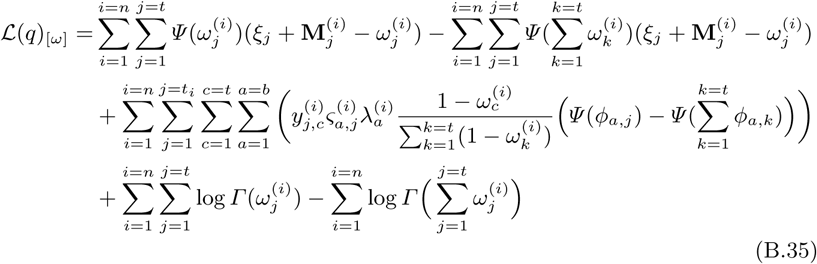

Taking derivatives w.r.t. 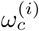 gives

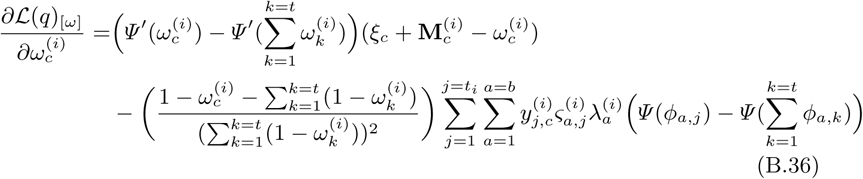

Setting it’s derivatives to zero does not lead to a closed-form solution, instead, we approximate 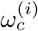 for each example *i* according to:

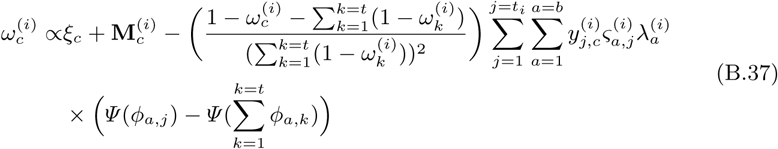

##### 6. Optimizing w.r.t

*λ*. Collecting the terms that contain *λ*, we obtain:

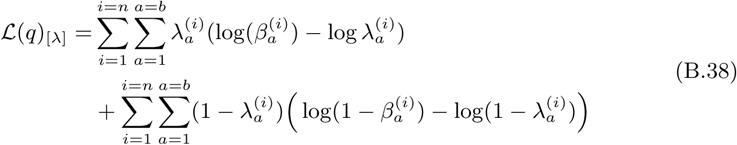

Taking derivatives w.r.t. 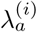, we obtain:

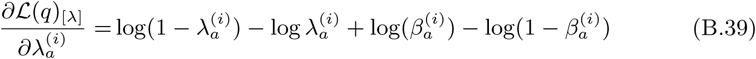

Equating the above formula to zero to obtain a maximum, we get the canonical parameterisation of the Bernoulli distribution:

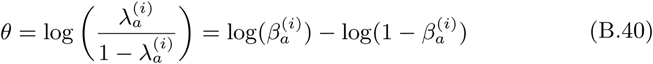

Therefore, we get the following updates:

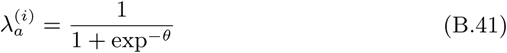

##### 7. Optimizing w.r.t *ϕ*

Finally, the optimal solution of the variational concept feature distribution *q*(Φ*a* | *ϕa*) for each concept *a* is obtained by isolating terms involved in the bound Eq. B.4:

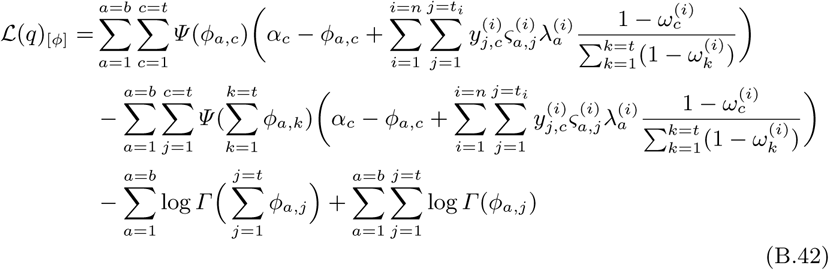

After taking derivatives w.r.t. *ϕa,c*, we obtain:

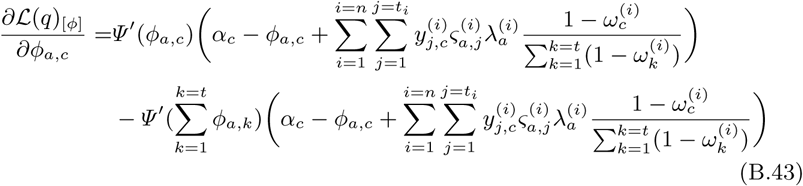

Equating the above formula to zero to obtain a maximum, we get:

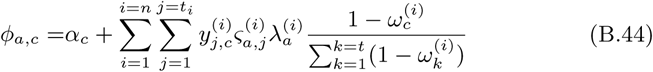

The variational inference algorithm samples a mini-batch from a collection, and use it to compute the local latent parameters in Eqs B.27, B.29, B.31, B.34, B.37, and B.41 until the evidence lower bound in Eq. B.4 converges. Then, the global variational parameter *ϕ* is updated using the posteriors (*β*, Λ, *η, z, Ω*) collected from the previous step in Eq. B.44, after being scaled according to the learning rate *τ* = (*s* + *l*)−*g*, where *s* is the current step, *l* ≥ 0 is the delay factor, and *g* ∈ (0.5, 1] is the forgetting rate. The variational inference process for SPREAT is summerized in Algorithm 7.

###### Algorithm 7: Stochastic variational inference for SPREAT

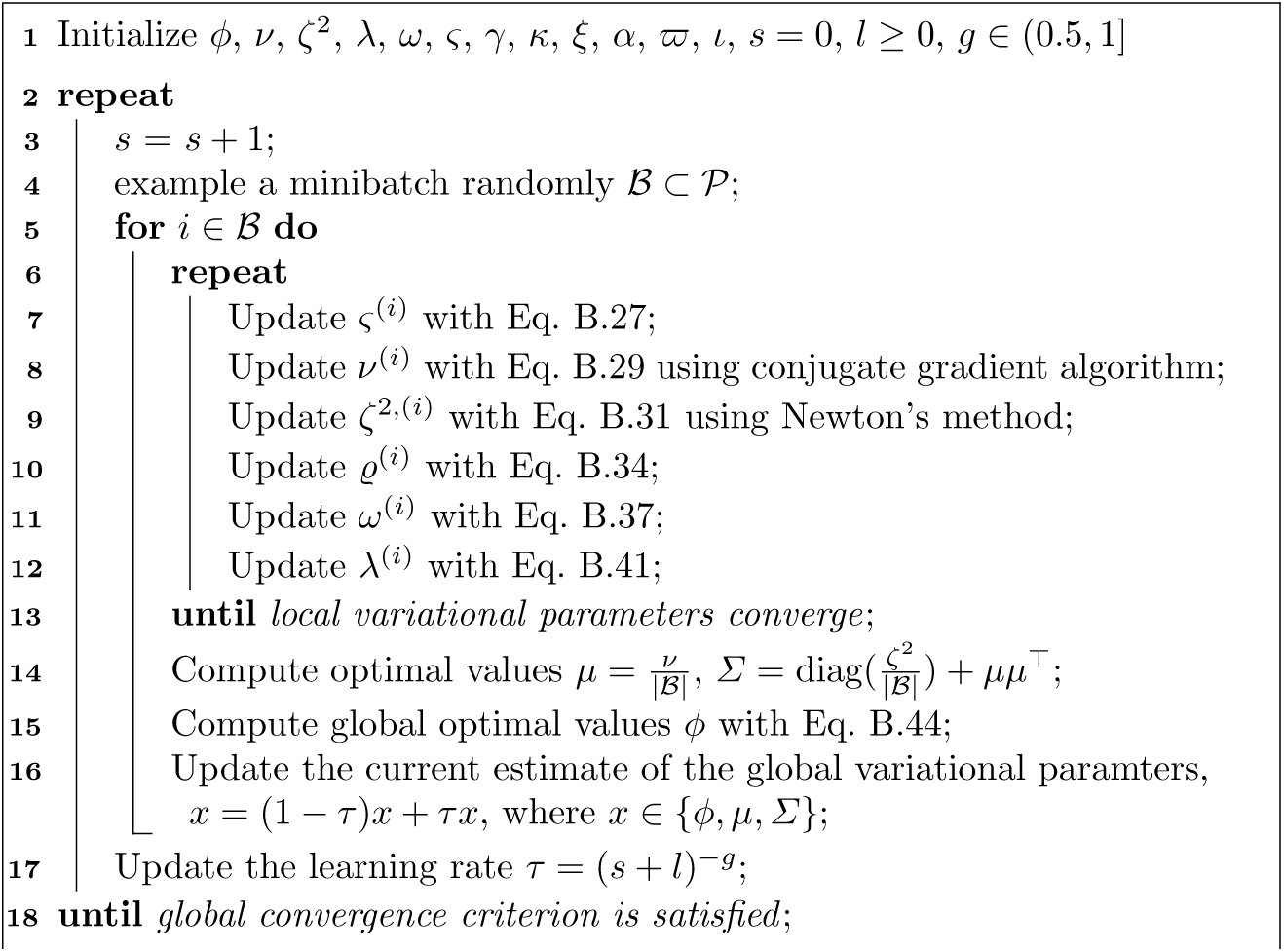

#### B.5 Posterior Predictive Distribution for SPREAT

The posterior predictive distribution is a useful and practical method to evaluate model’s fitness and to compare models without requiring to compute bounds of those models. This metric estimates the distribution of an unobserved value 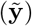 given the observed values (**Y***obs*) and parameters (*Θ* and **V**) that are trained on a held-out training set [31]. The predictive distribution for SPREAT is:

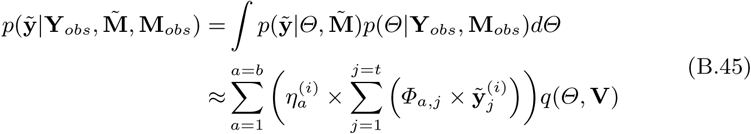

where *q*(*Θ*, **V**) corresponds to Eq. B.5 and trained on **Y***obs* and **M***obs*.

### C Experimental Setup

In this section, we describe the experimental settings and outline the materials used to evaluate the performance of reMap. The reMap and correlated models were written in Python v3 and depend on third party libraries (e.g. Numpy [49]). Unless otherwise specified all tests were conducted on a Linux server using 10 cores of Intel Xeon CPU E5-2650.

#### C.1 Description of Datasets

We used 11 simulated, organismal, and multi-organismal datasets to evaluate reMap’s grouping performance: i)- BioCyc v20.5 T2 & 3 [1], ii)- 6 T1 golden data that are composed of six databases (*EcoCyc (v21), HumanCyc (v19*.*5), AraCyc (v18*.*5), YeastCyc (v19*.*5), LeishCyc (v19*.*5)*, and *TrypanoCyc (v18*.*5)*), iii)- Symbiont genomes describing distributed metabolic pathways between *Moranella* (GenBank NC-015735) and *Tremblaya* (GenBank NC-015736) [15], iv)- Critical Assessment of Metagenome Interpretation (CAMI) initiative low complexity dataset [16], consisting of 40 genomes and is obtained from edwards.sdsu.edu/research/camichallenge-datasets/; v)- whole genome shotgun sequences from HOTS at 25m,

75m, 110m (sunlit) and 500m (dark) ocean depth intervals [17], and vi)- Synset- 2, a noisy corrupted training set [11]. The detailed characteristics of the datasets are summarized in Table 4. For each dataset 𝒮, we use |𝒮| and L(𝒮) to represent the number of instances and pathway labels, respectively. In addition, we also present some characteristics of the multi-label datasets, which are denoted as:

**Table 4:**
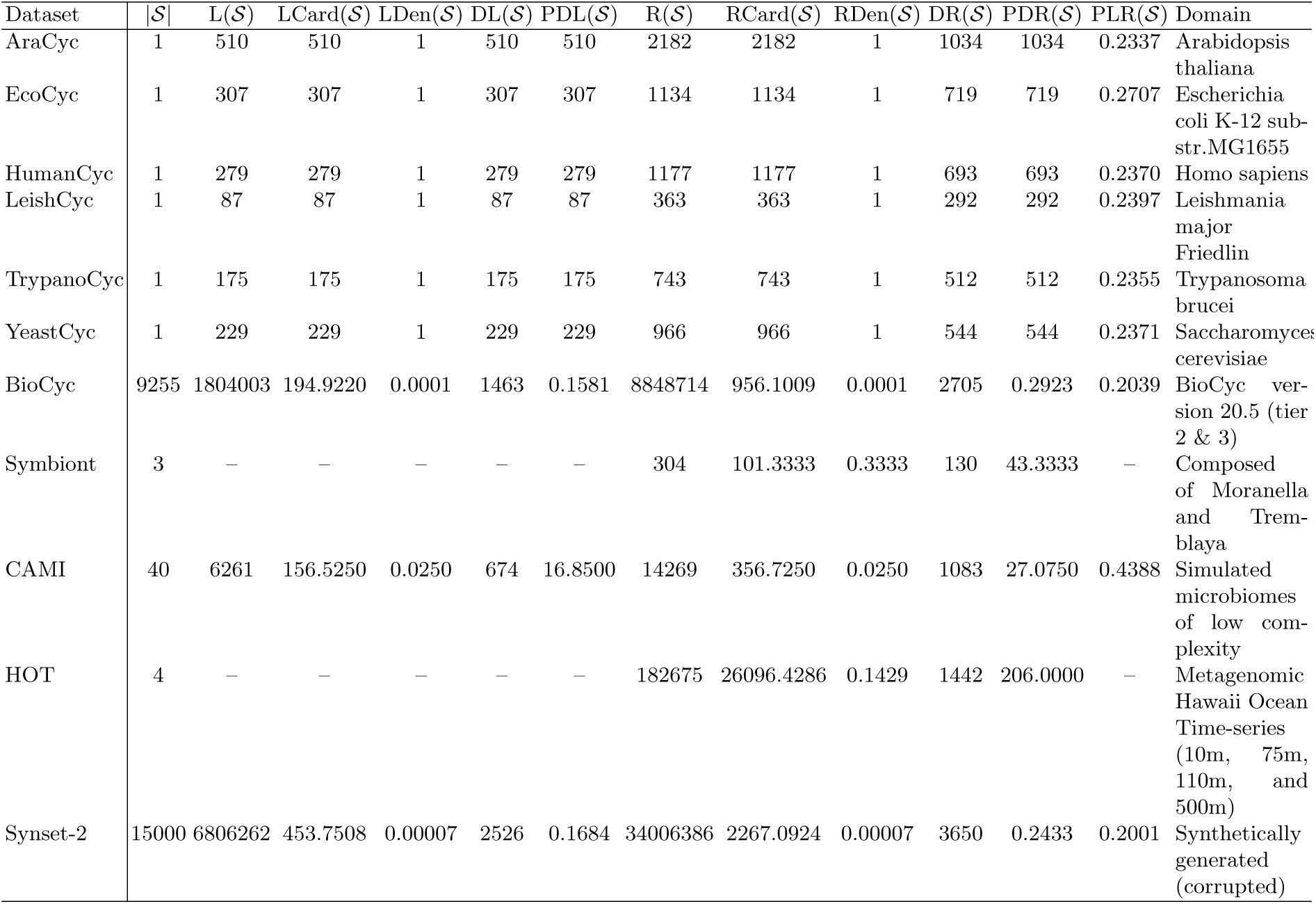
Characteristics of the experimental datasets. The notations |*𝒮*|, L(*𝒮*), LCard(*𝒮*), LDen(*𝒮*), DL(*𝒮*), and PDL(*𝒮*) represent number of instances, number of pathway labels, pathway labels cardinality, pathway labels density, distinct pathway labels, and proportion of distinct pathway labels for *𝒮*, respectively. The notations R(*𝒮*), RCard(*𝒮*), RDen(*𝒮*), DR(*𝒮*), and PDR(*𝒮*) have similar meanings as before but for the enzymatic reactions *E* in *𝒮*. PLR(*𝒮*) represents a ratio of L(*𝒮*) to R(*𝒮*). The last column denotes the domain of *𝒮*.

1. Label cardinality (LCard 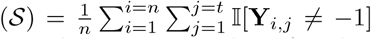), where 𝕀 is an indicator function. It denotes the average number of pathways in 𝒮.
2. Label density (LDen 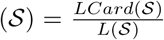). This is simply obtained through normalizing LCard(𝒮) by the number of total pathways in 𝒮.
3. Distinct pathway labels (DL(𝒮)). This notation indicates the number of distinct pathways in 𝒮.
4. Proportion of distinct pathway labels 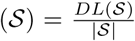. It represents the normalized version of DL(*𝒮*), and is obtained by dividing DL(.) with the number of instances in *𝒮*.

The notations R(𝒮), RCard(𝒮), RDen(𝒮), DR(𝒮), and PDR(𝒮) have similar meanings as before but for the enzymatic reactions ε in. Finally, PLR(𝒮) represents a ratio of L(𝒮) to R(𝒮). The preprocessed experimental datasets can be obtained from https://zenodo.org/record/3971534#.YX9dpWDMK3A.

#### C.2 Parameter Settings

We applied the following default configurations:

##### 1. reMap’s parameters

The parameters for the reMap model are configured as: the learning rate is *η* = 0.0001, the batch size is 30, the number of epochs is *τ* = 10, the group centroid hyperparameter is *α* = 16, the cutoff threshold for cosine similarity is *v* = 0.2, the cutoff decision threshold for groups is *β* = 0.3, the number of groups is *b* = 200, and the subsampled group size is *γ* = 50. For regularized hyperparameters *λ*1:5 and *κ*, we performed 10-fold cross-validation on a sample of BioCyc data (v20.5 T2 &3) and found the settings *λ*1:5 = 0.01 and *κ* = 0.01 to be the optimum.

##### 2. Correlated models parameters

The parameters for the three correlated models are configured as: the pathway distribution over concepts Φ are initialized using gamma distribution (with shape and scale parameters are fixed to 100 and 1*/*100, respectively), the forgetting rate is *g* = 0.9, the delay rate is *l* = 1, the batch size is 100, the number of epochs is 3, the number of concepts is *b* = 200, top *k* pathways is 100 (only for SOAP and SPREAT), the Dirichlet hyperparameters *α* and *ξ* are 0.0001, and the beta hyperparameters *γ* and *κ* are 2 and 3, respectively. The supplementary pathways **M** for BioCyc, CAMI, and golden T1 datasets are obtained using mlLGPR [11] trained on Synset-2. A schematic view of pathway frequency across datasets for BioCyc T2 & 3 and CAMI, along with their augmented pathways is depicted in Fig. 7.

**Fig. 7:**
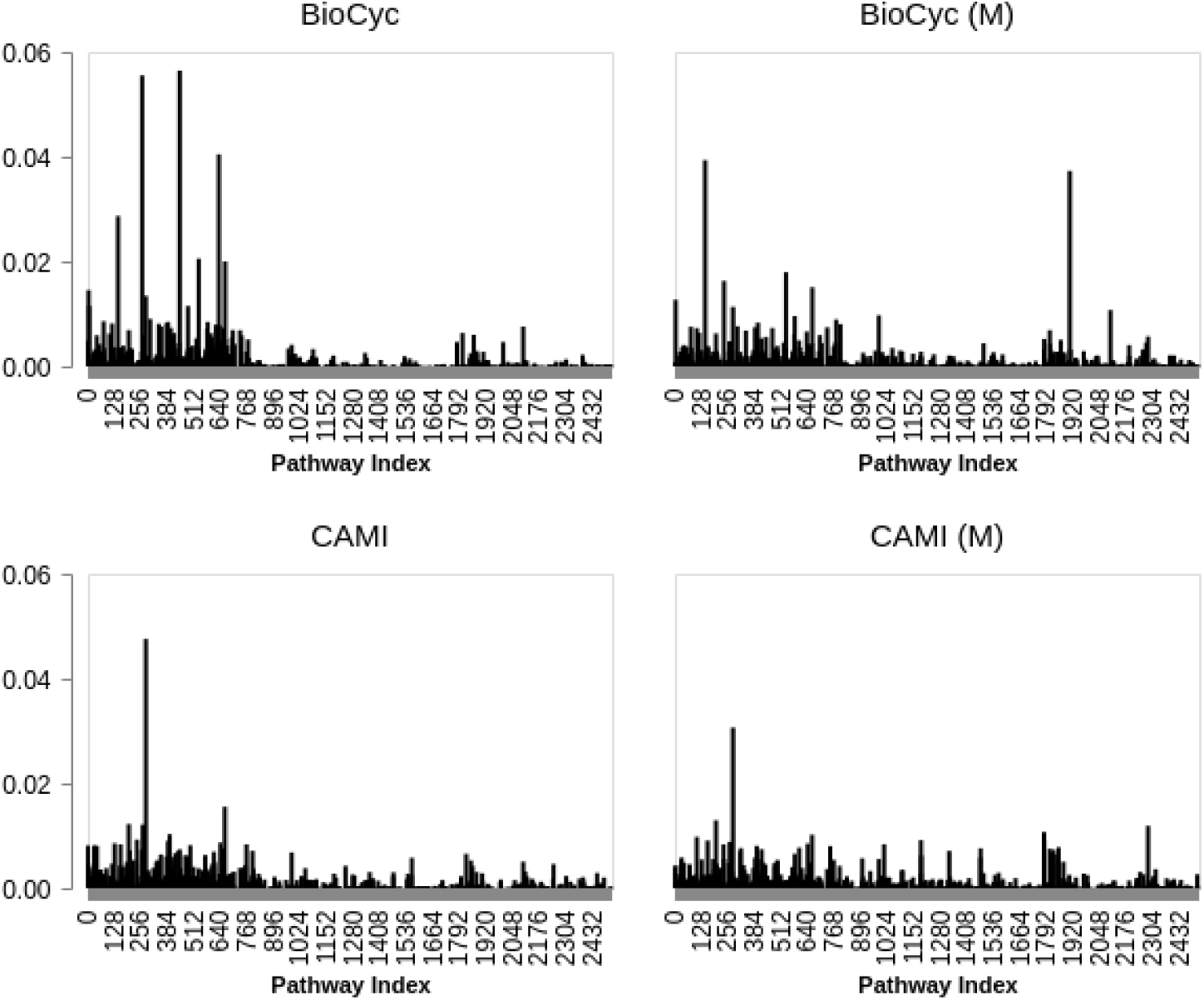
Pathway frequency (averaged on all examples) in BioCyc (v20.5 T2 &3) and CAMI data, and their background pathways, indicated by **M**.

##### 3. pathway2vec’s parameters

The parameters for the pathway2vec framework [10] are configured as: “crt” as the embedding method, the number of memorized domain is 3, the explore and the in-out hyperparameters are 0.55 and 0.84, respectively, the number of sampled path-instances is 100, the walk length is 100, the embedding dimension size is *m* = 128, the neighborhood size is 5, the size of negative examples is 5, and the used configuration of MetaCyc is “uec”, indicating trimmed links among ECs.

Both reMap and correlated models are trained using BioCyc (v20.5 T2 &3) collection. After obtaining groups *𝒮group*, we train leADS [14] using built-in “factorization” option that enables training pathway groups and pathways, simultaneously, for 10 epochs using “nPSP” as the acquisition function and “pref- voting” as the prediction strategy with cutoff threshold 0.5. For all the remaining hyperparameters in pathway2vec, correlated models, leADS, and mlLGPR [11], they are fixed to their default values.

### D Experimental Results

Two tests were performed to benchmark the performance of reMap including parameter sensitivity for correlated models and metabolic pathway prediction.

#### D.1 Sensitivity Analysis of Correlated Models

A fundamental challenge for the reMap pipeline is to acquire a good distribution of groups and pathways from correlated models for the purpose of relabeling. Following the common practice, here we examined various hyperparameters associated with correlated models. First, we compared the sensitivity of SOAP and SPREAT against CTM by incorporating the background pathways **M** while varying the number of groups according to *b* ∈ {50, 100, 150, 200, 300}. Next, we examined the “c2m” option for SOAP and SPREAT to show that these two models exhibit similar performances as CTM. Finally, we conducted sparsity analysis of group distribution by varying the cutoff threshold value according to *k* ∈ {50, 100, 150, 200, 300, 500}. For the comparative analysis, we applied CAMI as a test data to report the log predictive distribution (Section B.5), where a lower score entails higher generalization capability for the associated models.

While the log predictive scores for SOAP and SPREAT in Fig. 8a appears to be flat across group size, the CTM model projects a more realistic view where it’s performances are seen to be gaining by including more groups. For the former models, this phenomena is expected due to the effects of supplementary pathways. That is, both models are encouraged to learn more pathways from **M** because the average pathway size for an example in **M** is ≈ 500 whereas in BioCyc v20.5 T2 & 3 is ≈ 195. By excluding **M** (“c2m”), we observe that the log predictive distribution of SOAP and SPREAT are similar with that of CTM, as shown in Fig. 8b, which supports our previous discussion. From Figs 8a and 8b, it is evident that *b* = 200 represents the optimum group size with the average number of distinct pathways is ≈ 15. By fixing *b* = 200, we search for an optimum *k* value. As illustrated in Fig. 8c, both SOAP and SPREAT deteriorate their performances (*<*− 0.6) when *k >* 100. Taken together, we suggest the settings *b* ∈ ℤ[150,300] and *k* ∈ ℤ[50,100] to recover good pathway group and pathway distributions.

**Fig. 8:**
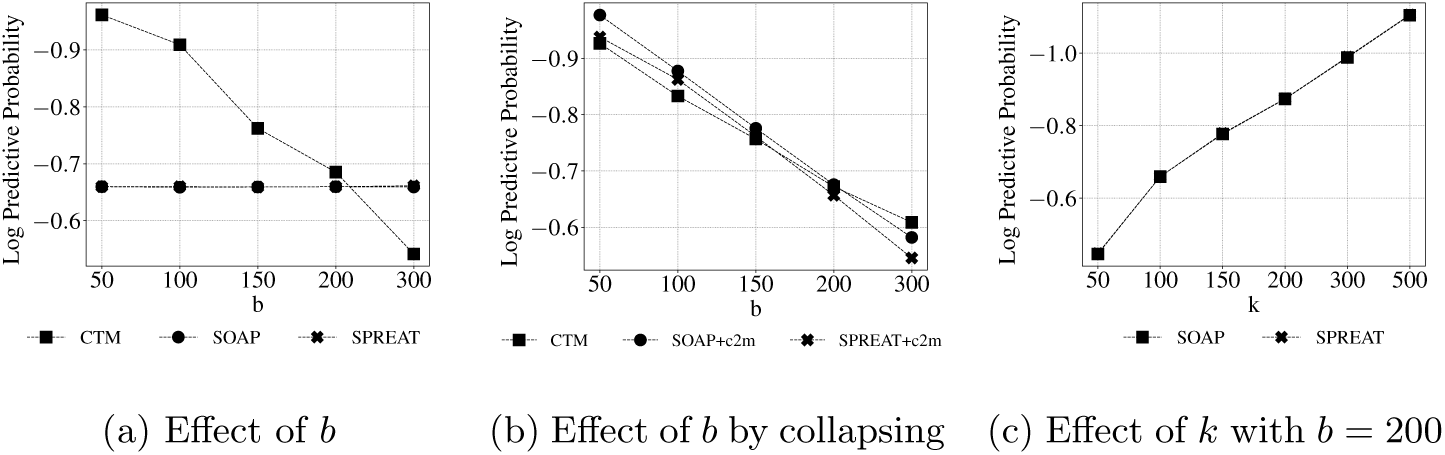
Log predictive distribution on CAMI data.

**Fig. 9:**
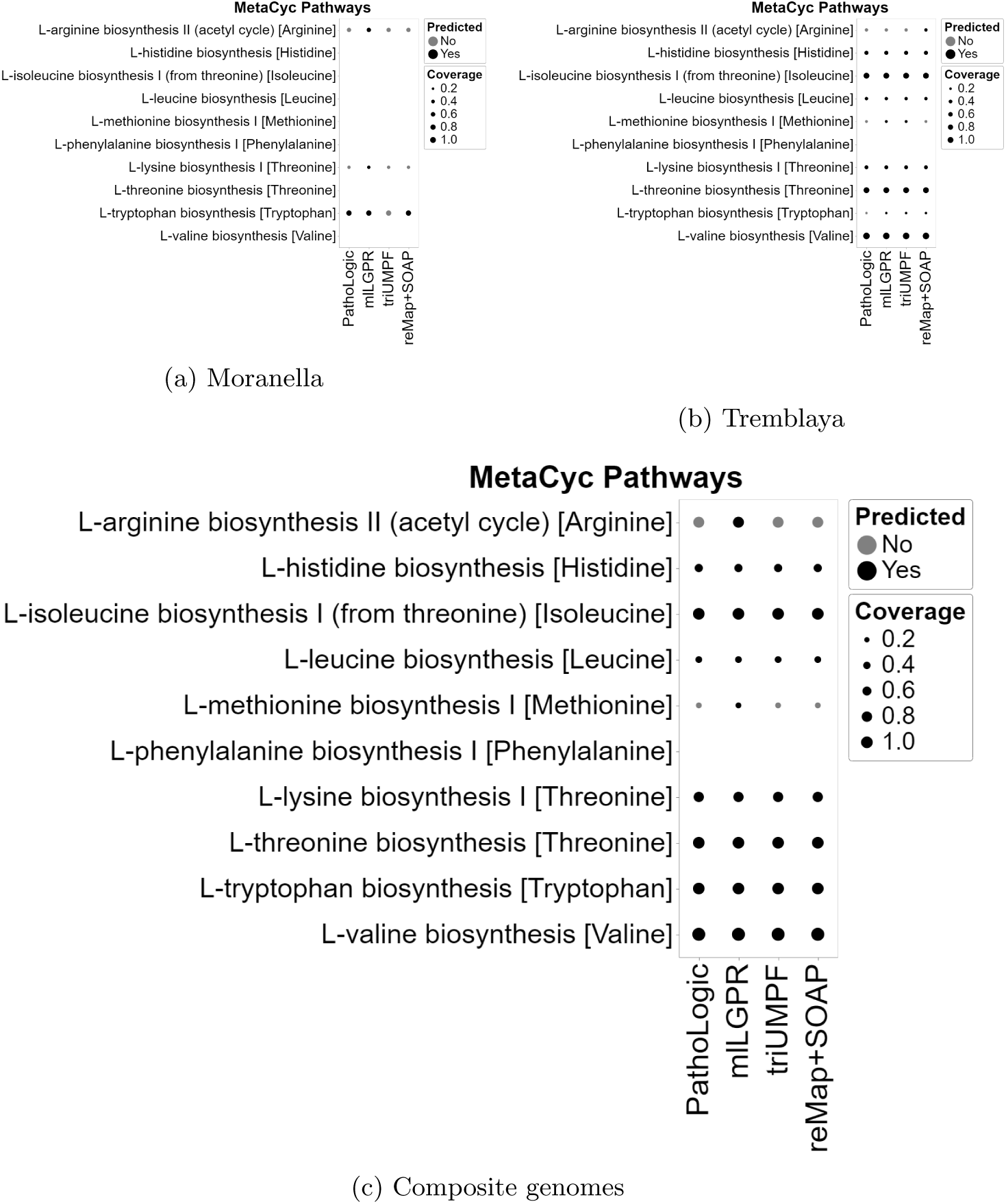
Comparative study of predicted pathways for symbiont data between PathoLogic, mlLGPR, triUMPF, and reMap+SOAP. Black circles indicate predicted pathways by the associated models while grey circles indicate pathways that were not recovered by models. The size of circles corresponds the pathway coverage information.

#### D.2 Accumulated History Probability Analysis

Table 5 shows 22 amino acid biosynthesis pathways with their 28 variants. Table 6 represents the selected 7 pathway groups that contain these amino acid pathways in their top 5 pathways for Escherichia coli K-12 MG1655 organism.

**Table 5:** 22 amino acid biosynthesis pathways and 28 pathway variants.

| Amino Acid | MetaCyc Pathway ID | MetaCyc Pathway Name |
| --- | --- | --- |
| Glycine | GLYSYN-PWY | glycine biosynthesis I |
| Alanine | ALANINE-VALINESYN-PWY | L-alanine biosynthesis I |
|  | ALANINE-SYN2-PWY | L-alanine biosynthesis II |
|  | PWY0-1021 | L-alanine biosynthesis III |
| Arginine | ARGSYN-PWY | L-arginine biosynthesis I (via L-ornithine) |
| Asparagine | ASPARAGINE-BIOSYNTHESIS | L-asparagine biosynthesis I |
|  | ASPARAGINESYN-PWY | L-asparagine biosynthesis II |
| Aspartate | ASPARTATESYN-PWY | L-aspartate biosynthesis |
| Chorismate | PWY-6163 | chorismate biosynthesis from 3-dehydroquinate |
|  | ARO-PWY | chorismate biosynthesis I |
| Cysteine | CYSTSYN-PWY | L-cysteine biosynthesis I |
|  | PWY-7870 | L-cysteine biosynthesis VII (from S-sulfo-L-cysteine) |
| Glutamate | GLUTSYN-PWY | L-glutamate biosynthesis I |
|  | GLUTSYNIII-PWY | L-glutamate biosynthesis III |
| Glutamine | GLNSYN-PWY | L-glutamine biosynthesis I |
| Histidine | HISTSYN-PWY | L-histidine biosynthesis |
| Isoleucine | ILEUSYN-PWY | L-isoleucine biosynthesis I (from threonine) |
| Leucine | LEUSYN-PWY | L-leucine biosynthesis |
| Lysine | DAPLYSINESYN-PWY | L-lysine biosynthesis I |
| Methionine | HOMOSER-METSYN-PWY | L-methionine biosynthesis I |
| Phenylalanine | PHESYN | L-phenylalanine biosynthesis I |
| Proline | PROSYN-PWY | L-proline biosynthesis I |
| Selenocysteine | PWY0-901 | L-selenocysteine biosynthesis I (bacteria) |
| Serine | SERSYN-PWY | L-serine biosynthesis |
| Threonine | HOMOSER-THRESYN-PWY | L-threonine biosynthesis |
| Tryptophan | TRPSYN-PWY | L-tryptophan biosynthesis |
| Tyrosine | TYRSYN | L-tyrosine biosynthesis I |
| Valine | VALSYN-PWY | L-valine biosynthesis |

**Table 6:**
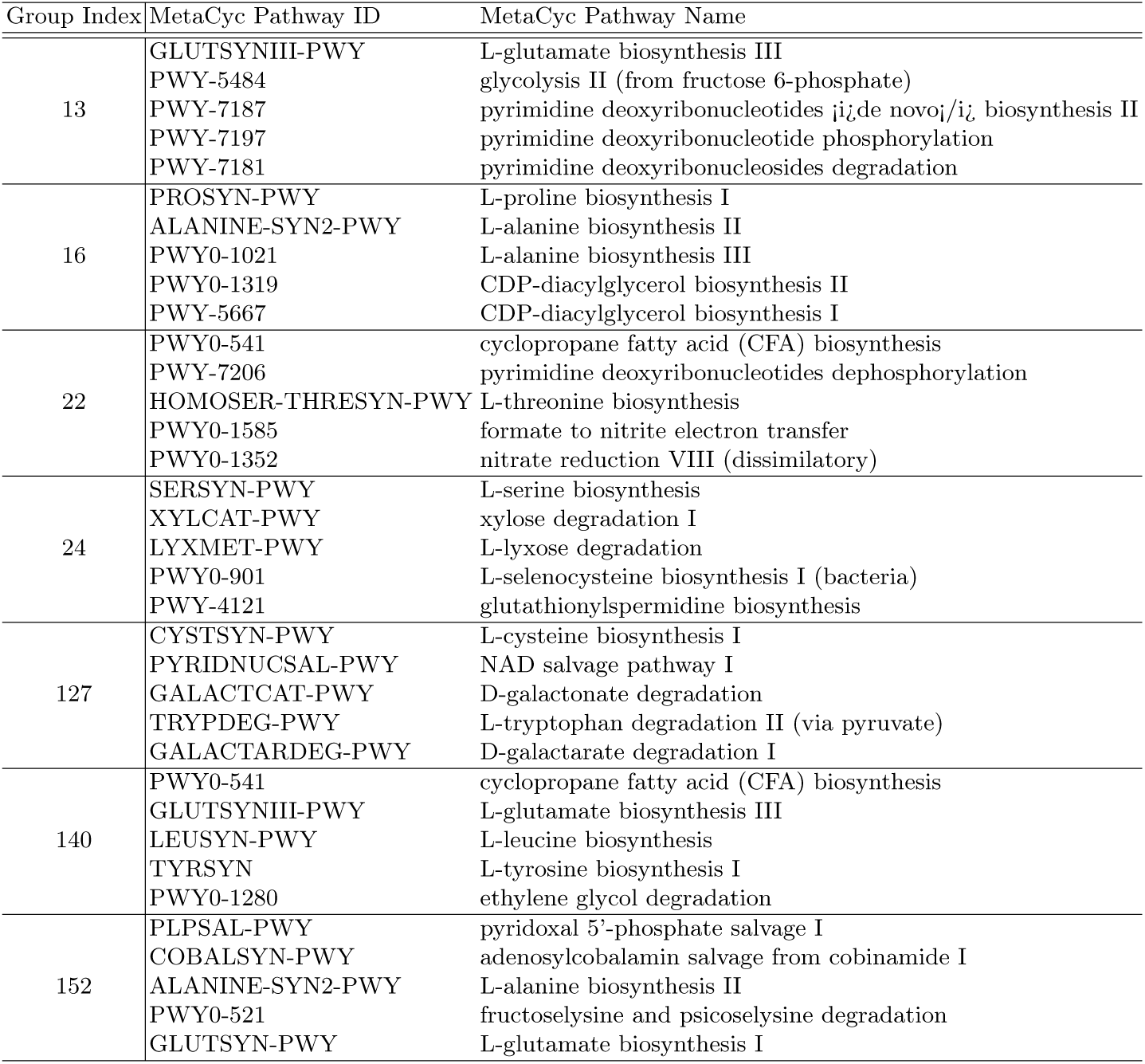
7 pathway groups containing 22 amino acids and their top 5 pathways for Escherichia coli K-12 MG1655.

| Group Index | MetaCyc Pathway ID | MetaCyc Pathway Name |
| --- | --- | --- |
| 13 | GLUTSYNIII-PWY | L-glutamate biosynthesis III |
|  | PWY-5484 | glycolysis II (from fructose 6-phosphate) |
| | PWY-7187 | pyrimidine deoxyribonucleotides $\mu_i$de novo/$i_L$ biosynthesis II |
|  | PWY-7197 | pyrimidine deoxyribonucleotide phosphorylation |
|  | PWY-7181 | pyrimidine deoxyribonucleosides degradation |
| 16 | PROSYN-PWY | L-proline biosynthesis I |
|  | ALANINE-SYN2-PWY | L-alanine biosynthesis II |
|  | PWY0-1021 | L-alanine biosynthesis III |
|  | PWY0-1319 | CDP-diacylglycerol biosynthesis II |
|  | PWY-5667 | CDP-diacylglycerol biosynthesis I |
| 22 | PWY0-541 | cyclopropane fatty acid (CFA) biosynthesis |
|  | PWY-7206 | pyrimidine deoxyribonucleotides dephosphorylation |
|  | HOMOSER-THRESYN-PWY | L-threonine biosynthesis |
|  | PWY0-1585 | formate to nitrite electron transfer |
|  | PWY0-1352 | nitrate reduction VIII (dissimilatory) |
| 24 | SERSYN-PWY | L-serine biosynthesis |
|  | XYLCAT-PWY | xylose degradation I |
|  | LYXMET-PWY | L-lyxose degradation |
|  | PWY0-901 | L-selenocysteine biosynthesis I (bacteria) |
|  | PWY-4121 | glutathionylspermidine biosynthesis |
| 127 | CYSTSYN-PWY | L-cysteine biosynthesis I |
|  | PYRIDNUCSAL-PWY | NAD salvage pathway I |
|  | GALACTCAT-PWY | D-galactonate degradation |
|  | TRYPDEG-PWY | L-tryptophan degradation II (via pyruvate) |
|  | GALACTARDEG-PWY | D-galactarate degradation I |
| 140 | PWY0-541 | cyclopropane fatty acid (CFA) biosynthesis |
|  | GLUTSYNIII-PWY | L-glutamate biosynthesis III |
|  | LEUSYN-PWY | L-leucine biosynthesis |
|  | TYRSYN | L-tyrosine biosynthesis I |
|  | PWY0-1280 | ethylene glycol degradation |
| 152 | PLPSAL-PWY | pyridoxal 5'-phosphate salvage I |
|  | COBALSYN-PWY | adenosylcobalamin salvage from cobinamide I |
|  | ALANINE-SYN2-PWY | L-alanine biosynthesis II |
|  | PWY0-521 | fructoselysine and psicoselysine degradation |
|  | GLUTSYN-PWY | L-glutamate biosynthesis I |

#### D.3 Metabolic Pathway Prediction

Here, groups obtained from all correlated modules are used for the pathway prediction task. For this, we trained leADS using the configuration discussed in Section C.2. The results are reported on golden T1 and CAMI data using four evaluation metrics: *Hamming loss, average precision, average recall*, and *average F1 score*. We also studied reMap’s performance on Symbiont and HOTS data. For comparative analysis, four pathway prediction algorithms are used: i)- MinPath v1.2 [20], ii)- PathoLogic v21 [7], iii)- mlLGPR (elastic net with enzymatic reaction and pathway evidence features) [11], and iv)- triUMPF [13].

Table 7 shows that reMap+SOAP outperforms triUMPF on five T1 golden data (excluding LeishCyc) with regard to average recall and average F1 scores where numbers in boldface represent the best performance score in each column while the underlined text indicates the best performance among correlated models. For the remaining correlated models, their sensitivity scores are higher than triUMPF with the exception to EcoCyc and AraCyc. Similar results are observed for Symbiont, CAMI, and HOTS (Figs 10, 11, 12, and 13) data. In summary, this experiment demonstrates that pathway group based approach, in particular reMap+SOAP, improves pathway predictions.

**Table 7:** Predictive performance of each comparing algorithm on 6 benchmark datasets. For each performance metric, ‘↓’ indicates the smaller score is better while ‘↑’ indicates the higher score is better. Bold text suggests the best performance in each column.

| Methods | Hamming Loss ↓ |  |  |  |  |  |
| --- | --- | --- | --- | --- | --- | --- |
|  | EcoCyc | HumanCyc | AraCyc | YeastCyc | LeishCyc | TrypanoCyc |
| PathoLogic | 0.0610 | 0.0633 | 0.1188 | **0.0424** | **0.0368** | **0.0424** |
| MinPath | 0.2257 | 0.2530 | 0.3266 | 0.2482 | 0.1615 | 0.2561 |
| mlLGPR | 0.0804 | 0.0633 | **0.1069** | 0.0550 | 0.0380 | 0.0590 |
| triUMPF | 0.0435 | 0.0954 | 0.1560 | 0.0649 | 0.0443 | 0.0776 |
| reMap+SOAP | **0.0392** | **0.0400** | 0.1714 | 0.0934 | 0.0772 | <u>0.0479</u> |
| reMap+SPREATE | 0.0519 | 0.0827 | 0.1489 | <u>0.0748</u> | 0.0629 | 0.0503 |
| reMap+CTM | 0.0558 | 0.0835 | <u>0.1425</u> | 0.0804 | 0.0622 | 0.0503 |
| reMap+SOAP+c2m | 0.0590 | 0.0780 | 0.1457 | 0.0772 | 0.0614 | 0.0534 |
| reMap+SPREATE+c2m | 0.0542 | 0.0796 | 0.1520 | 0.0772 | <u>0.0598</u> | 0.0558 |
| Methods | Average Precision Score ↑ |  |  |  |  |  |
|  | EcoCyc | HumanCyc | AraCyc | YeastCyc | LeishCyc | TrypanoCyc |
| PathoLogic | 0.7230 | 0.6695 | 0.7011 | 0.7194 | **0.4803** | 0.5480 |
| MinPath | 0.3490 | 0.3004 | 0.3806 | 0.2675 | 0.1758 | 0.2129 |
| mlLGPR | 0.6187 | 0.6686 | 0.7372 | 0.6480 | 0.4731 | 0.5455 |
| triUMPF | 0.8662 | 0.6080 | 0.7377 | **0.7273** | 0.4161 | 0.4561 |
| reMap+SOAP | 0.8611 | **0.7871** | 0.6215 | 0.4851 | 0.2805 | 0.5985 |
| reMap+SPREATE | **0.9400** | 0.6750 | 0.8350 | 0.6000 | 0.3200 | **0.6200** |
| reMap+CTM | 0.9150 | 0.6700 | **0.8750** | 0.5650 | 0.3250 | **0.6200** |
| reMap+SOAP+c2m | 0.8950 | 0.7050 | 0.8550 | 0.5850 | 0.3300 | 0.6000 |
| reMap+SPREATE+c2m | 0.9250 | 0.6950 | 0.8150 | 0.5850 | <u>0.3400</u> | 0.5850 |
| Methods | Average Recall Score ↑ |  |  |  |  |  |
|  | EcoCyc | HumanCyc | AraCyc | YeastCyc | LeishCyc | TrypanoCyc |
| PathoLogic | 0.8078 | 0.8423 | 0.7176 | 0.8734 | 0.8391 | 0.7829 |
| MinPath | **0.9902** | **0.9713** | **0.9843** | **1.0000** | **1.0000** | **1.0000** |
| mlLGPR | 0.8827 | 0.8459 | 0.7314 | 0.8603 | 0.9080 | 0.8914 |
| triUMPF | 0.7590 | 0.3835 | 0.3529 | 0.3319 | 0.7126 | 0.6229 |
| reMap+SOAP | <u>0.8078</u> | <u>0.8746</u> | <u>0.3863</u> | 0.4978 | <u>0.7931</u> | <u>0.9371</u> |
| reMap+SPREATE | 0.6124 | 0.4839 | 0.3275 | <u>0.5240</u> | 0.7356 | 0.7086 |
| reMap+CTM | 0.5961 | 0.4803 | 0.3431 | 0.4934 | 0.7471 | 0.7086 |
| reMap+SOAP+c2m | 0.5831 | 0.5054 | 0.3353 | 0.5109 | 0.7586 | 0.6857 |
| reMap+SPREATE+c2m | 0.6026 | 0.4982 | 0.3196 | 0.5109 | 0.7816 | 0.6686 |
| Methods | Average F1 Score ↑ |  |  |  |  |  |
|  | EcoCyc | HumanCyc | AraCyc | YeastCyc | LeishCyc | TrypanoCyc |
| PathoLogic | 0.7631 | 0.7460 | 0.7093 | **0.7890** | 0.6109 | 0.6447 |
| MinPath | 0.5161 | 0.4589 | 0.5489 | 0.4221 | 0.2990 | 0.3511 |
| mlLGPR | 0.7275 | 0.7468 | **0.7343** | 0.7392 | **0.6220** | 0.6768 |
| triUMPF | 0.8090 | 0.4703 | 0.4775 | 0.4735 | 0.5254 | 0.5266 |
| reMap+SOAP | **0.8336** | **0.8285** | 0.4764 | 0.4914 | 0.4144 | **0.7305** |
| reMap+SPREATE | 0.7416 | 0.5637 | 0.4704 | <u>0.5594</u> | 0.4460 | 0.6613 |
| reMap+CTM | 0.7219 | 0.5595 | <u>0.4930</u> | 0.5268 | 0.4530 | 0.6613 |
| reMap+SOAP+c2m | 0.7061 | 0.5887 | 0.4817 | 0.5455 | 0.4599 | 0.6400 |
| reMap+SPREATE+c2m | 0.7298 | 0.5804 | 0.4592 | 0.5455 | <u>0.4739</u> | 0.6240 |

**Fig. 10:**
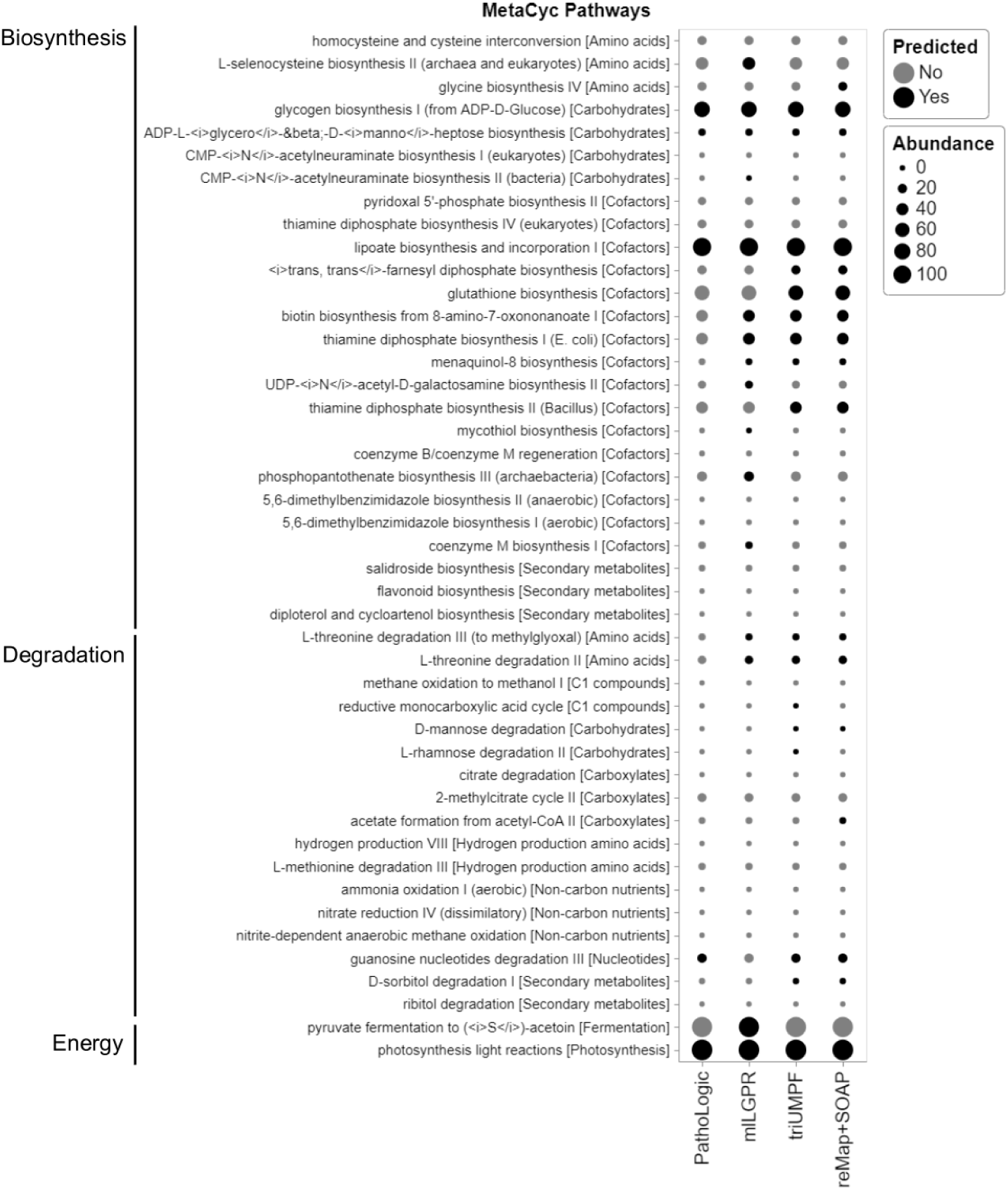
Comparative study of predicted pathways for HOTS 25m dataset between PathoLogic, mlLGPR, triUMPF, and reMap+SOAP. Black circles indicate predicted pathways by the associated models while grey circles indicate pathways that were not recovered by models. The size of circles corresponds the pathway abundance information.

**Fig. 11:**
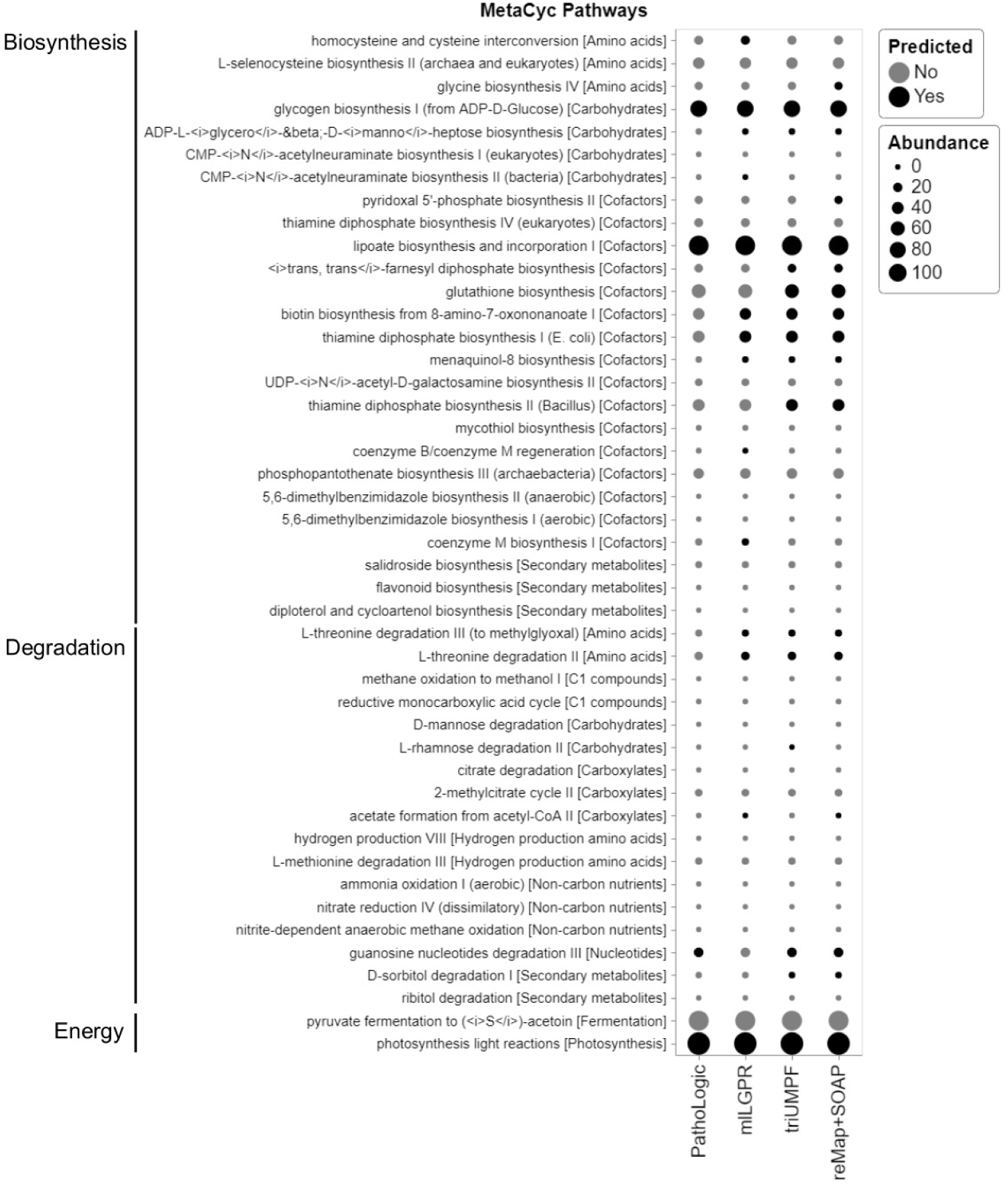
Comparative study of predicted pathways for HOTS 75m dataset between PathoLogic, mlLGPR, triUMPF, and reMap+SOAP. Black circles indicate predicted pathways by the associated models while grey circles indicate pathways that were not recovered by models. The size of circles corresponds the pathway abundance information.

**Fig. 12:**
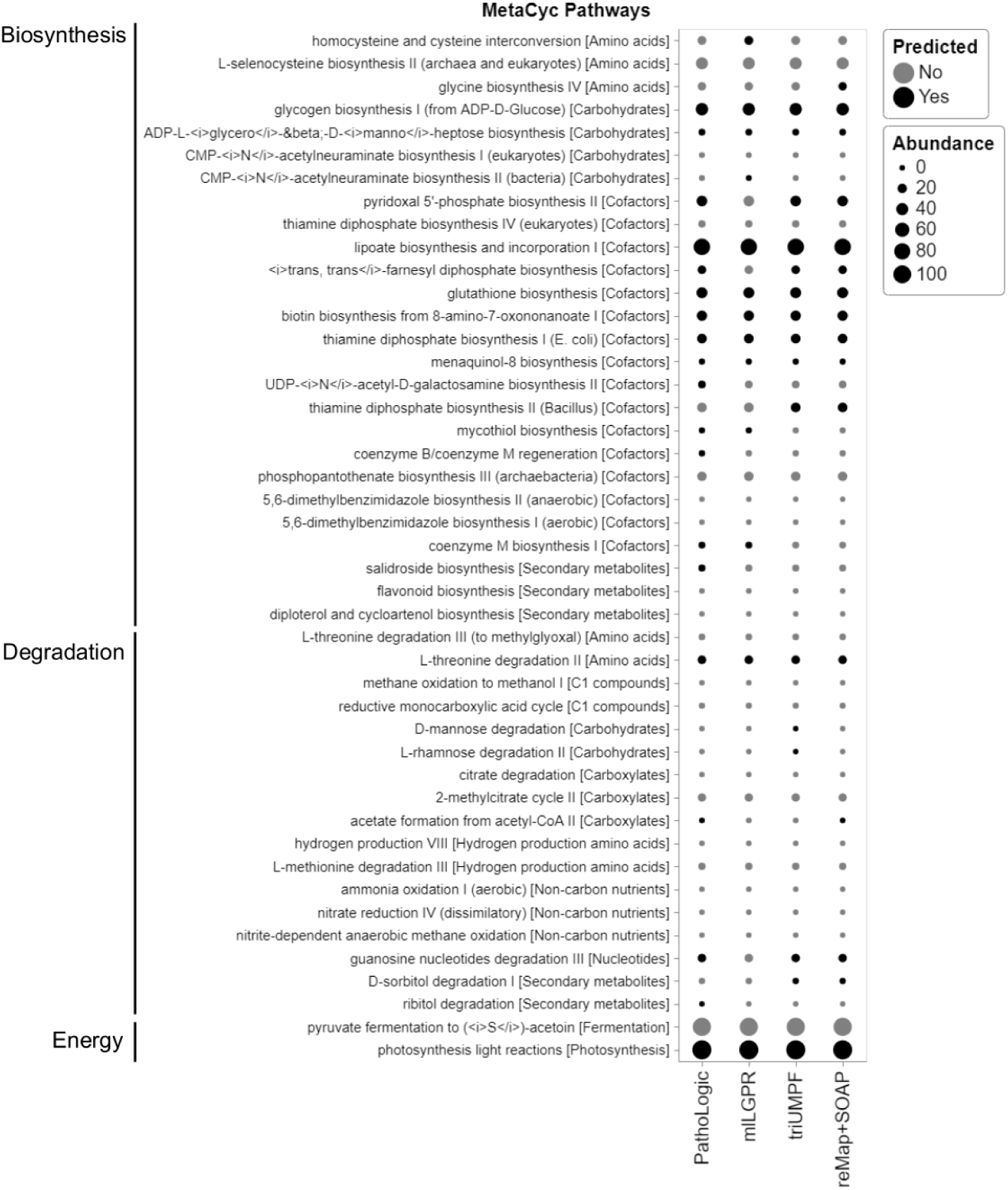
Comparative study of predicted pathways for HOTS 110m dataset between PathoLogic, mlLGPR, triUMPF, and reMap+SOAP. Black circles indicate predicted pathways by the associated models while grey circles indicate pathways that were not recovered by models. The size of circles corresponds the pathway abundance information.

**Fig. 13:**
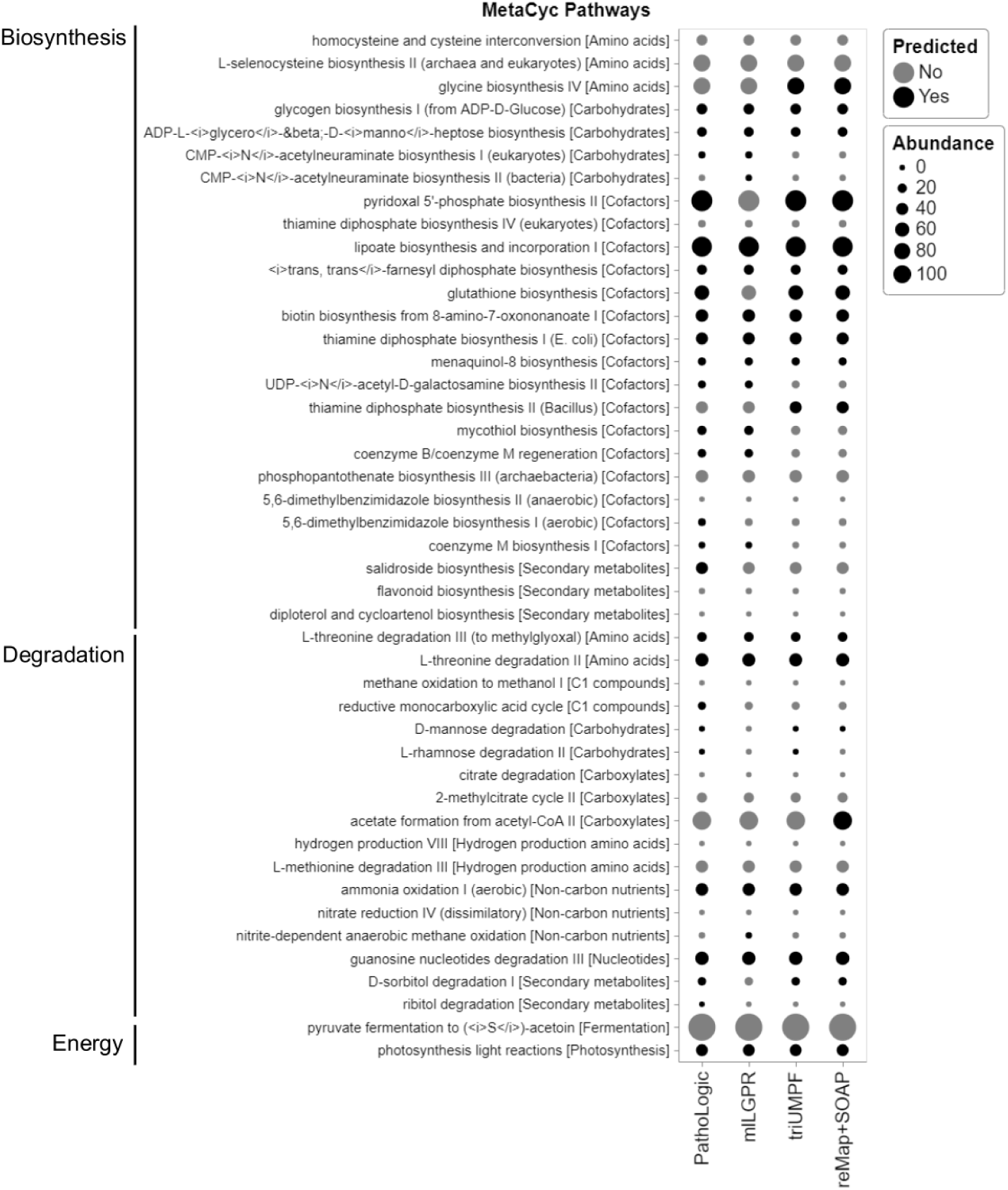
Comparative study of predicted pathways for HOTS 500m dataset between PathoLogic, mlLGPR, triUMPF, and reMap+SOAP. Black circles indicate predicted pathways by the associated models while grey circles indicate pathways that were not recovered by models. The size of circles corresponds the pathway abundance information.

